# Human populations with low survival at advanced ages and postponed fertility reduce long-term growth in high inflation environments

**DOI:** 10.1101/2024.07.31.606043

**Authors:** Rahul Mondal, José Manuel Aburto, Rebecca Sear, Shripad Tuljapurkar, Uday Shankar Mishra, Roberto Salguero-Gómez

## Abstract

Temporal variability in inflation can lead to important fluctuations in the long-term growth rate of human populations via their differential impacts on vital rates like survival and fertility. However, historically, demographic studies have overlooked this time-dependent relationship. Here, we test whether human populations have higher stochastic population growth rates when exposed to lower levels of inflation. We also examine if lower survival rates at older ages (>60 years) and fertility rates at the later reproductive years (>30 years) among populations exposed to higher inflation rates determine their expected lower long-term growth rate compared to those exposed to lower inflation rates. To explore the impact of variability in inflation on vital rates response, we develop a quantitative pipeline with four steps, and parameterise it with high-resolution economic and demographic data across 76 countries from 1971-2021. The four steps are (1) defining treatment groups based on levels of trend inflation (creeping inflation (0-3%), walking inflation (3-10%), galloping inflation (10-50%), and hyperinflation (>50%)) among which the stochastic population growth rates will be compared; (2) constructing matrix population models for each environmental state under every treatment. The environmental states for each treatment are defined on the basis of the duration of inflation (*e.g*., 0, 2, 4, six years or above); (3) estimating the stochastic population growth rate for each treatment by considering a Markovian environment dictated by the long-term frequency (*f*) and temporal autocorrelation (*p*) of the treatment; and (4) decomposing the differences in the population growth rate between treatments into contributions from environmental variability and vital rate differences between environments to test how vital rates impact on population growth under varying environmental scenarios. In agreement with our hypothesis, we find that the stochastic population growth rate at lower levels of inflation is systematically higher than that at a higher level of inflation at all stationary frequencies and temporal autocorrelation of the inflation environment. Moreover, the disadvantage in survival at older ages (>60 years) and fertility at ages >30 years led to the lower stochastic growth rate among populations exposed to higher level of inflation such as walking inflation compared to lower level of inflation, such as creeping inflation. Our framework explicitly links human population performance and inflation environment by describing nonlinear feedback between inflation, human survival, fertility, population growth, and its age structure. We discuss the potential of our approach to study the life-history strategies and population dynamics of a wide range of drivers of environmental variability.

## Introduction

Temporal variability in inflation can lead to large fluctuations in long-term human population growth rate. This effect can occur through the differential effects of the former on the vital rates of survival and fertility between environments. Economic factors, including inflation, are important exogenous drivers of human survival (Cutler et al 2006; Deaton 2003; Preston 1975), fertility (Becker 1960, 1992; Easterlin 1975), and ultimately of population growth rates (He 2018). However, historically, demographic studies have overlooked how time-dependent variation in economic factors shape demographic characteristics (but see Lee 1985; Schultz 1985; Tuljapurkar 1989). To understand and forecast population responses to variable economic environments, we need to consider time-varying random vital rates (Coale 1957; Cohen 1977a; Lee 1974; Sykes 1969). Thus, the examination of population dynamics in variable environments requires the random vital rates defined within varying economic environments (Lee and Tuljapurkar 1994; Tuljapurkar 2005). These economic dynamics and volatility are best described by inflation levels (*e.g*., Ascari and Sbordone 2014; Stock and Watson 2007, 2016) due to their ties with monetary policy and external shocks (Ascari & Sbordone 2014). High and variable inflation coupled with high labour market and lifetime uncertainty may reduce investment in old-age survival (Edwards 2013; Ehrlich 2000; Ehrlich and Becker 1972; Garcia-Sanchez and Pierrard 2023) and postponed fertility (Sobotka et al. 2011), reducing long-term population growth. In contrast, increasing fertility at early ages of mothers can act as a strategy against low juvenile survival at a highly variable inflation environment (Kalemli-Ozcan 2003, 2008; LeGrand and Sandberg 2006; Yakita 2001). Hence, identifying ‘inflation environments’ can provide a robust base to understand the interplay of time-varying vital rates and environmental states in determining long-term population dynamics.

How vital rates respond to variability in a given inflation environment can be a useful descriptor of population dynamics (Tuljapurkar 1982a, 1982b). Indeed, an extensive body of literature exists describing population dynamics as the population growth rate and its dependence on the temporal variation in vital rates or characteristics of a Markov Process (Caswell 2001, 2009, 2019; Cohen 1977a, 1977b, 1979; Tuljapurkar 1982a, 1982b; Tuljapurkar and Orzack 1980). The impact of variability in vital rates depends on the sensitivity of the population growth rate to changes in the average vital rates in each environment (Tuljapurkar 1982a, 1982b, 1989). In variable environments, vital rates with large sensitivities are viewed as major determinants of population growth (Tuljapurkar 1982a). In this context, Tuljapurkar’s small noise approximation (1982a, 1982b) shows how sensitivity analysis for average vital rates reveals significant features of population dynamics in fluctuating environments. Therefore, sensitivities of population growth rate to vital rates in a Markovian environment enable us to quantify the impact of the variability in inflation environments on population performance.

Defining environments based on inflation levels and examining their impacts on population dynamics poses several challenges. First, inflation itself has nonlinear time-varying dynamics (Ascari and Sbordone 2014, Stock and Watson 2016). Second, multiple sources of noise exist regarding inflation data, and the nature of said sources of noise is time-varying (Stock and Watson 2007, 2016). Thus, the accurate estimation of the impacts of inflation on population dynamics requires filtering out these noises (Atkeson and Ohanian 2001; Cogley and Sargent 2015; Nelson and Schwert 1977; Stock and Watson 2007, 2016). Third, senescence in human survival and fertility is slow-moving and difficult to reverse (Bobeica et al. 2017; Juselius and Takats 2018) as the response of survival and fertility rates to changing environments has a unique pattern and a lag effect due to bio-social constraints (Aburto et al. 2020; Colchero et al. 2016). Therefore, the research community is yet to successfully isolate the effects of inflation on human population dynamics from other secular trends (*e.g*., technological innovation [Coccia 2014; Dosi and Nelson 2010; Peretto 1998; Weinberger et al. 2017], cultural changes [Ghirlanda et al. 2010; Richerson et al. 2009; Shennan 2000]).

Research explicitly accounting for inflation dynamics and isolating the impact of variability in the inflation environment on population dynamics is particularly valuable for human demography (Juselius and Takats 2018). These studies will provide answers to the essential question of how economic volatility shapes demographic response and determine the asymptotic population dynamics. Here, we aim to delineate the inflation-population dynamics by deviating from classical economic and demographic studies on three areas. First, we incorporate the theory of random rates in variable environments in reference to both inflation and population dynamics (Ascari and Sbordone 2014; Cohen 1977a; Stock and Watson 2016; Tuljapurkar 1989). Second, we explicitly consider inflation trends, as this key component largely dominates the dynamics of inflation (Stock and Watson 2007). Finally, our study considers the lag effect of vital rates response to inflation and accounts for the impact of other secular trends like cultural and technological changes by the time-varying component of the framework. Thus, our approach takes a key step towards isolating the effect of inflation dynamics on population variability by filtering out the effect of unexplained factors.

Here, we introduce a quantitative framework to examine how variability in the inflation environment and the response of vital rates translate into fluctuations in population growth rates. We apply this framework to the World Bank’s Global Database on inflation to define the economic environment and the United Nations World Population Prospects 2022 version (WPP, hereafter; United Nations 2022) to construct matrix population models to test the following hypotheses: (H1) Human populations have higher stochastic population growth rates when exposed to lower levels of inflation, regardless of frequency and temporal autocorrelation in the inflation environment; (H2) lower survival rates at older ages (>60 years) and fertility rate at late reproductive ages (30 years) among population exposed to higher inflation rates would determine their lower long-term growth rate compared to those exposed to lower inflation rates. To test both hypotheses, we implement a stochastic Life Table Response Experiment (sLTRE, hereafter; Caswell 2010, 2019) to decompose the impact of temporal variability in the inflation environment and vital rates response to the environment on differences in stochastic population growth at different inflation environments. Our inflation-dependent demography framework is an essential step towards linking economic stochasticity to macro-demographic change.

## Materials and Methods

To test the hypotheses on the impacts of temporal variability in inflation environment on long-term population growth rate, we developed a quantitative pipeline. This pipeline relies on the *a priori* classification of populations according to country, inflation level (*treatment*, hereafter), and duration in the said inflation level (*environmental state*, hereafter). We first define the treatment groups based on levels of trend inflation among which the stochastic population growth rate will be compared (Figure 1A). Then, we construct projection matrices for each environmental state under every treatment (Fig. 1B). In the next section, we calculate stochastic growth rate for each treatment by considering a Markovian environment (Fig. 1E). Finally, we decompose the differences in the stochastic population growth rate between treatments into 1) contributions from environmental variability and 2) vital rates differences between environments to test the hypothesis on impact of vital rates on population growth at varying environmental dynamics, characterised by stationary frequency and temporal autocorrelation (Fig. 1F).

**Fig. 1.**
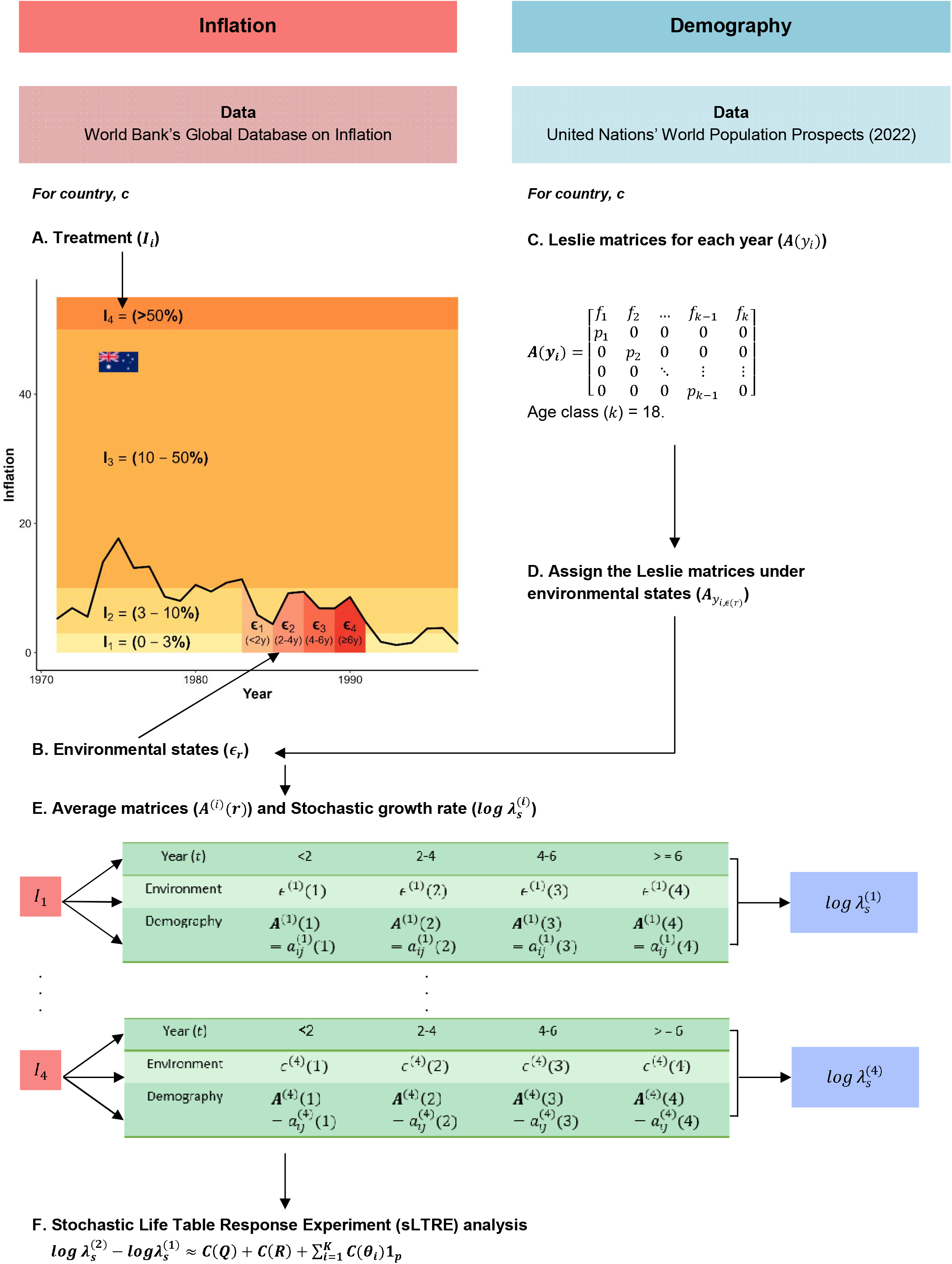
Design of the stochastic Life Table Response Experiment (sLTRE) analysis to decompose the difference in stochastic growth rate between treatments into contributions from differences in environmental dynamics and vital rates under each environment. Using trend inflation data from the World Bank’s Global Database on Inflation (A), we defined four treatments (*I*_*m*_), *i.e*., *I*_1_, *I*_2_, *I*_3_, and *I*_4_, based on the level of inflation. The Global Database provides trend inflation data across 76 countries from 1971-2022. Therefore, for each country (*c*), each year (*y*_*i*_) was assigned to one of the four treatments (*I*_*m*_) corresponding to the level of inflation. (B) We defined environmental states (*ϵ*_*r*_) for each treatment, i.e., ϵ_1_, *ϵ*_2_, *ϵ*_3_, and *ϵ*_4_, based on the duration of treatment (*I*_*m*_) in the country, *c*. We assigned years in each treatment,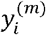, to an environmental state (*ϵ*_*r*_), based on the corresponding duration of treatment (*I*). For instance, in year *y*_*i*_, the country *c* went to treatment *I*_*m*_ for the first time, the year (*y*_*i*_) will be assigned to environmental state *ϵ*_1_ for treatment *I*_*m*_, i.e., 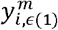, or if in year *y*_*i*_ the country *c* completes three years in that treatment *I*_*m*_, the year (*y*_*i*_) will be assigned to *ϵ*_2_ under *I*_*m*_ for that country, *c*, i.e, 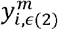. (C) We used the demographic data (survival and fertility rates) from the United Nations World Population Prospects, 2022, to construct Leslie matrices for each year, *A*(*y*_*i*_), from 1971-2022 for the 76 countries. To link inflation with demography, (D) we assigned the matrices (*A*(*y*_*i*_)) under one of the environmental states (*ϵ*_*r*_) by matching their corresponding year (*y*_*i*_) with the years under environmental states under each treatment 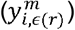 *i.e*., 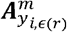 To focus on the duration of treatment (E), we constructed a set of average matrices, *A*^*(m)*^*(ϵ*_*r*_), one for each environmental state (*E*_*r*_). Each average matrix *A*^*(m)*^(*ϵ*_*r*_) is the element-by-element arithmetic mean of all the year-wise Leslie matrices 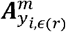 corresponding to that environmental state *ϵ*_*r*_. We then calculated the long-term stochastic growth rate for each treatment, 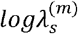 The growth rates were calculated by Markov chain simulation with *T* =1000 iterations. (F) Finally, we decomposed the differences in stochastic growth rate between (i) *I*_1_ and *I*_2_, (ii) *I*_2_ and *I*_3_, and (iii) *I*_3_ and *I*_4_ into the contributions from differences in stationary frequency, *C*(***Q***), temporal autocorrelation pattern, *C*(***R***), and vital rates in each environmental state, *C*(***θ***_*i*_) between the treatments, *I*_*m*_.

### Inflation Data

To explore the impact of inflation on the long-term growth of human populations, we extracted inflation data from the World Bank’s Global Database on Inflation (Ha et al. 2023). This comprehensive database includes six measures of inflation. These measures are headline, food, energy, core consumer price index inflation, producer price index inflation, and gross domestic product deflator. These measures are available at three frequencies (monthly, quarterly, and annual) across 209 countries between 1970 and 2022. Ha et al. (2023) provides the estimates of trend inflation, using a univariate component stochastic volatility model, for 76 countries from 1971-2022 on a quarterly frequency in the Global Database on Inflation. These estimates of trend inflation filter out time-varying noise and accounts for their volatility in inflation rates (Ascari and Sbordone 2014; stock and Watson 2007). Thus, these estimates can be used to describe economic dynamics (Ascari and Sbordone 2014). Our analyses of the impact of trend inflation on human population dynamics use the trend inflation data for the 76 countries for which estimates are available in Ha et al. (2023). Variability in vital rates and representativity of the examined countries is detailed in the Appendix and Discussion.

### Matrix Population Models

Examining how populations respond to a variable inflation environment requires estimates of their stochastic growth rate. Here, we opted to use age-based models (so-called Leslie matrices; Leslie 1945, 1948) corresponding to different inflation rates. To that end, we consider a single population whose individuals are in one of the *k* age classes, and where time is measured in discrete intervals. Survival and fertility rates are explicitly contained in a Leslie matrix (Caswell 1978). Population growth rate and change in structure from one time-step to the next is given by ***n***(*t*+1)=**A**|***n***(*t*)|, where ***n****(t)* is the population vector that describes the number of individuals of a given age (*i* =1, …, *k*) in the population at time *t*, and where **A** is the Leslie matrix. The Leslie matrix is an *n*×*n* matrix with non-zero elements possible only in the first row (*i.e*., fecundity) and the sub-diagonal (*i.e*., survival). Conventional Leslie matrices recognize *n* age groups before the highest age of female reproduction, 50, yielding a 10 × 10 matrix (Geramita and Pullman 1978; Kim and Schoen 1997; Schoen and Kim 1991). Our projection matrix **A**, however, considers 18 age classes from ages 0 to 85+, producing **A**_18×18_ matrices.

We constructed Leslie matrices for multiple human populations using age-specific survival and fertility rates from the United Nations’ World Population Prospects (WPP, hereafter; United Nations, 2022). This resource contains vital rate estimates for 237 nations or regions from 1950 to the present, supported by analyses of historical demographic trends. Data from 1,758 national population censuses, vital registration systems, and 2,890 nationally representative sample surveys conducted between 1950 and 2022 are considered in this most recent evaluation. We adopted the survival probabilities from the abridged life table from 1971-2021 from the WPP 2022 data and used them as the sub-diagonal entries of the Leslie matrix **A** for each of the 76 target countries. To calculate fertility for the age classes in the first row of **A**, we used the age-specific fertility rate and the sex ratio at birth from WPP 2022. We calculated fertility as

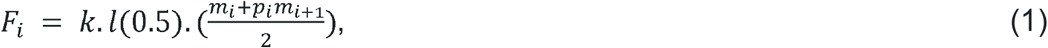

where *k* is the proportion of female birth calculated using the female sex ratio at birth (Preston et al. 2002), *l(0.5)* the probability that an average individual survives half of the projection interval, *m*_*i*_ the age-specific fertility rate, and *p*_*i*_ the survival rate of mothers. These Leslie matrices **A**(*y*_*i*_), corresponding to inflation rates for each country-year are next used to analyze the impact of inflation on human population dynamics (Fig. 1D).

### Treatments and Environments

To examine differences in stochastic population growth rate at different levels of inflation, we categorised projection matrices into treatments based on levels of inflation. The estimates of trend inflation obtained from the World Bank’s Global Database on Inflation (Ha et al 2022) ranged between -10.49% (*e.g*., Myanmar in 1973) and 738.77% (*e.g*., Argentina in 1985). A higher level of trend inflation tends to decrease average economic output, to unanchor inflation expectations, raise economic volatility, and reduce welfare (Ascari and Sbordone 2014). However, policy makers tend to target a higher inflation rate to reduce the risk of hitting the Zero Lower Bound (ZLB) constraints, where nominal interest rate is at or near zero, thus hindering economic growth (Ascari and Sbordone 2014). An inflation target of ∼2% is robust to a wide range of calibrations of hitting ZLB (Coibion et al. 2012; Fischers 1996; Summers 1991). Thus, in our comparative analysis, we consider an inflation level of 0-3% as the benchmark (*I*_0_: control). To define the treatments, we adopted the following established classification (Frisch, 1977; Lim and Sek, 2015; Machlup, 2020; Wolozin, 1959):, *I*_1_ : creeping inflation (0-3%), *I*_2_: walking inflation (3-10%), *I*_3_: galloping inflation (10-50%); and, *I*_4_: hyperinflation (>50%). Next, we assigned each of the Leslie matrices, **A**(*y*_*i*_),from 1971 to 2022 for each of our 76 countries into the treatment group .*I*_*m*_(*m* =1,…,4).

To quantify the environmental variability of each treatment,, *I*_*m*_, we defined environmental states. Briefly, these environmental states. ϵ_*r*_(*r*=1,…,4,) were assigned to each inflation treatment level, *I*_m_ based on the duration at said, *I*m for each country (Figure 1). As the age pattern of human survival and fertility does not change rapidly over time (Wachter 2003) due to bio-social constraints (Aburto et al. 2020; Colchero et al 2016), are typically characterised by time lags (Juselius and Takats 2018), and rarely reverse (Bobeica et al. 2017), we defined .*ϵr* based on the duration of, *I*m, as opposed to the levels of inflation. Here, the environments are divided into four states:. *ϵ*1 (<2 yrs), .ϵ_2_ (2-4 yrs), .ϵ3(4-6 yrs) and .ϵ_4_ (>6 yrs) (Fig. 1B). Thus, for each country, we categorised the Leslie matrices 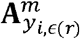 under each inflation treatment and environmental states within said treatment. However, to focus on the duration of, *I*_*m*_, we calculated the averaged matrices,, **A**^(^*m*^)^,.(ϵ_1_),…,,**A**^(^*m*^)^,. (*ϵ*_4_), (Fig. 1E) by calculating the element-by-element arithmetic mean of all year matrices, 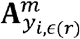 corresponding to one environmental state, .*ϵ*_*r*_. This approach resulted in a single matrix population model for each environmental state within each of the inflation treatments experience by each of our 76 examined countries (see Fig. 1E).

Therefore, in each treatment for every country, there are *r* possible environmental states and a matrix of vital rates for each environmental states,, **A**^(^*m*^)^,.(*ϵ*_*r*_), (The assumption of a finite number of environmental states can be relaxed for serially independent environments). The temporal sequence of environments is an ergodic aperiodic Markov process, meaning that the probability distribution of every sequence of environmental states eventually ‘forgets’ its past and converges to a stationary distribution (Caswell 2001; Tuljapurkar 1982a, 1989).

### Stochastic LTRE Analysis

Finally, to examine the impact of variable inflation environment on stochastic population growth rate 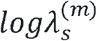, we performed a stochastic Life Table Response Experiment (sLTRE; Caswell 2010, 2019) analysis. LTREs were first introduced in stage-structured demography by Caswell (1989), and are often used in ecology and human demography to decompose effects of density (Sibly 1999; Oli et al. 2001), land use change (Jacquemyn and Brys 2008), nutrition (Cooch et al. 2001; Oli and Dobsoon 2001) fire (Caswell 2010; Caswell and Kaye 2001) and variable environment (Caswell 2019) on the asymptotic growth rates from deterministic matrix population models (Caswell 1989, 2000). However, this approach has also been applied to decomposing the impact of multiple drivers on stochastic population growth rates (Caswell 2010, 2019; Caswell and Kaye 2001; Davison et al. 2010), transient dynamics (Koons et al. 2016), and even life expectancy (Horiuchi et al. 2008; Pollard 1988) or measures of lifespan inequality (Van Raalte and Caswell 2013).

Here, we develop and apply an sLTRE analysis to decompose the difference in stochastic growth rate, 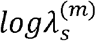 between treatments, *I*_*m*_ into impacts from variable inflation environment. The sLTRE approach adopted in this study is a key improvement in studying the effect of inflation on population dynamics for three important reasons. First, we account for the time-varying dynamics of inflation by defining the treatments based on levels of ‘trend’ inflation (Stock and Watson 2007). Second, we consider the lag effect of vital rates responses to variable inflation environment by defining environmental states as the duration of said inflation levels (Juselius and Takats 2018). Finally, the model isolates the effect of inflation from secular trends of unexplained variables (technology and cultural changes) as the latter are already adjusted by the time component in the time-varying environment in the model (Foreman et al. 2018; Lee and Carter 1992).

### Stochastic Population Growth Rate

To test (H1) whether the 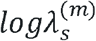 is higher at a lower inflation level, we calculated, 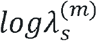 for each treatment at a varying inflation environment. With a Leslie matrix, **A**(*t*) generated by a stochastic environment that produces a set of vital rates for each environmental state, the asymptotic long-term growth rate with probability 1 is obtained as

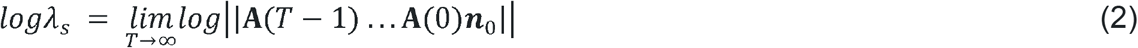

(Caswell 2019; Cohen 1976; Tuljapurkar 1990; Tuljapurkar and Orzack 1980). The abundance vector .***n***_0_ in equation 2 can be any fixed, arbitrary non-negative vector with ||n|| = 1, by the Furstenberg-Kersten theorem (Caswell 2001). This population growth rate is the stochastic analogue of the population growth rate in constant environments.

### Stochastic Inflation Environment

To test (H2) whether lower old-age survival and late-age fertility leads to lower 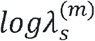 at a higher inflation level, we performed an sLTRE decomposition. The sLTRE decomposes the differences in 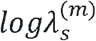 between treatments into contribution from two components: (1) differences in environmental dynamics, and (2) differences in vital rates within each environmental state. To do so, we generated a four-state Markov chain for the environmental states,. *ϵ*_*i*_ (Fig. 1B). The Markov chain transition matrix, **P**, determines the environmental dynamics where the probability distribution of the environmental state at time *t*+1 depends only on the state at time *t, p*_ij_ = **P**[*u*(*t*+1)=*I* |*u*(*t*)=*j*, . The resulting transition matrix is

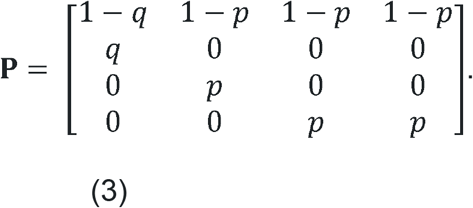

The entries of the transition matrix **P** are calculated from the frequency (*f*) and temporal autocorrelation (ρ) of the treatmeants,*I*_l_ and, *I*_2_, as follows

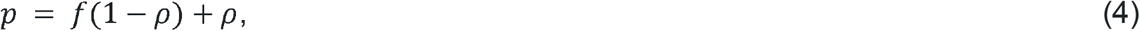

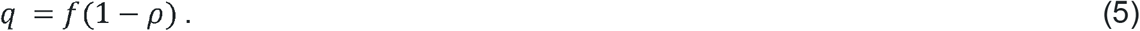

If ρ< 0,*f* must satisfy

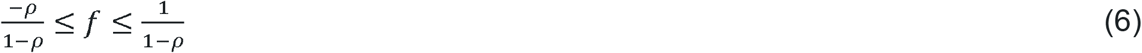

to keep probabilities bounded between 0 and 1(Caswell 2001; Caswell and Kaye 2001).

The matrices **P**^(l)^ and **P**^(2)^ may differ in their stationary distribution, pattern of autocorrelation, or both (Caswell 2001, 2019). These stationary distributions are described by the right eigenvector π, corresponding to the dominant eigenvalue of **P**. Therefore, to separate the contribution of long-term frequency (given by stationary distribution) and autocorrelation, we constructed a Markov chain with the same stationary distribution, π as **P**, but where successive environmental states are independent and identically distributed (iid), and hence have no temporal autocorrelation (Caswell 2001, 2019). This chain has the transition matrix

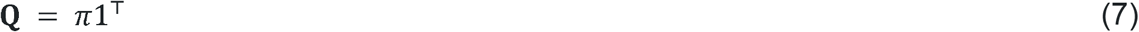

where **1** is a vector of ones. However, autocorrelations cannot have an arbitrarily large effect on population dynamics. Firstly, the elements of the vital rate matrices {A_*i*_} must obey evolutionary and biomechanical constraints (Tuljapurkar 1982a). Secondly, for any given set {A_*i*_}, there are limits to population dynamics irrespective of temporal pattern (Tuljapurkar 1982a).

### Environment-Specific Sensitivities

To calculate the contributions of vital rates to the difference in 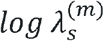 between, *I*_*m*_ we computed environment-specific sensitivities of vital rates to 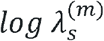 To do so, we computed the derivatives of log*λ*s with respect to each vector. θ_*i*_ in. *θ* (entries of the Leslie matrix,**A**) in each state of environments. The formula for computing the environment-specific sensitivities was given by Caswell (2005) and Horvitz et al. (2006).

To make the *logλ*_*s*_ as a function of **P** and ***θ***, we generated a stochastic sequence of *T* environments by Markov chain simulation of **P** and **Q** using the *markovchain* package in R (Spedicato et al. 2016), where *T* is a large number. Subsequently, we used the sequence to generate a sequence of Leslie matrices **A**_0_,**A**_1_,…,**A**_*T™*1_, and calculated the stochastic growth rate as

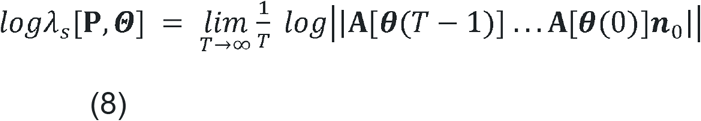

where θ(*t*) is the parameter vector created by the environmental state .*u*(*t*) (Caswell 2001, 2019). Then, we calculated the derivatives of *logλ*_*s*_ with respect to vital rates in environment *i* as

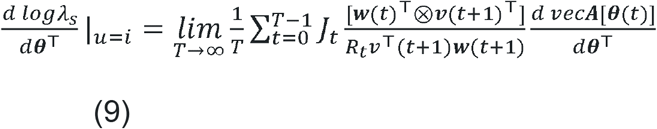

(Caswell 2001, 2019). Here, ***W***(*t*) is the right eigenvector describing the age distribution that is obtained by projecting forward in time, while, ***v***(*t*) is the left eigenvector that defines the reproductive value, computed by backward projection (Caswell 2001).Likewise, *R*_*t*_ is the growth of total population size from, *t* to, *t*+1. To make the sensitivity environment dependent, we incorporated the indicator variable,

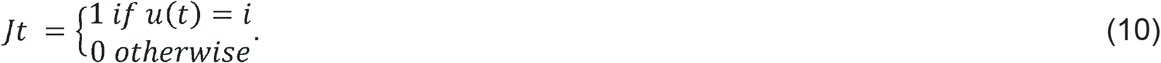

The last component, 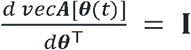, I where is the identity matrix, as the parameters *θ* are the elements of the Leslie matrix, *A* (Caswell, 2019).

### sLTRE Decomposition

Finally, to examine (H2) the impact of vital rates response to variable inflation environment on growth rate, we performed the sLTRE decomposition of .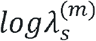 differences. We decomposed the differences in the stochastic growth rate of treatment 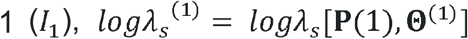 and treatment 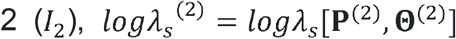 into contributions from differences in stationary environmental frequencies (*C*(**Q**)), autocorrelation pattern (*C*(**R**)), and vital rates in each environmental state (*C*(***θ***_1_),…,*C*(***θ***_*K*_)). As .*logλ*_*s*_ is computed by simulation, it cannot be differentiated with respect to (Caswell 2019; Steinsaltz et al. 2011). Thus, we used the Kitagawa decomposition (Kitagawa 1955) for calculating the environmental dynamics contributions as

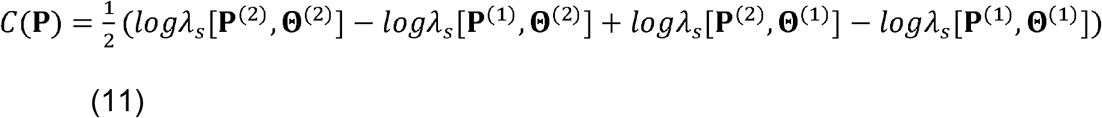

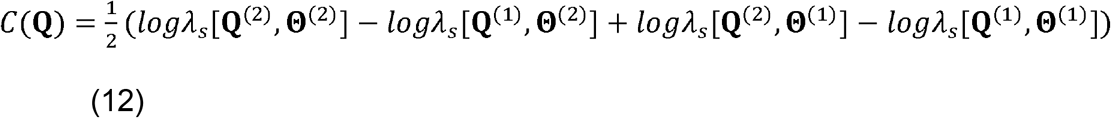

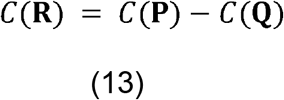

where each of C(**P**), C(**Q**), and C(**R**) is a scalar (Caswell, 2010, 2019). We expressed the contributions of vital rates in differences using the Kitagawa-Keyfitz method as

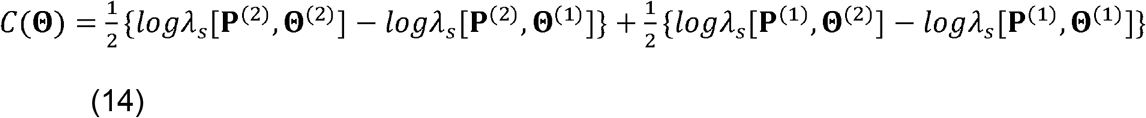

Where,*C*(Θ) is a scalar, obtained by summing the contributions of differences in all vital rates responses at all environmental states. We further decomposed each term in Equation 14 into contributions from the vital rates in each environment using environment-specific sensitivities as

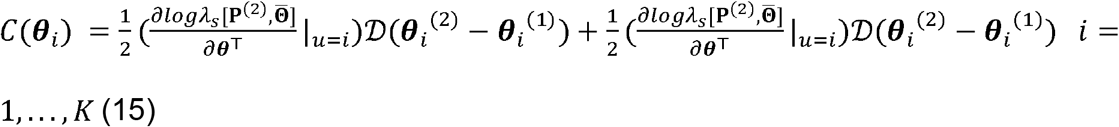

with derivatives evaluated the mean of vital rates between two treatments (*I*_l_ and*I*_2_),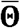 Thus, the matrix, *C*(***θ***_i_) is a vector whose elements give the contributions from differences in each vital rate from the environment. *ϵ*_*i*_ to differences in *log λ*_*s*_ (Caswell 2010, 2019).

To test the aforementioned hypotheses, we performed the sLTRE under two scenarios: hypothetical and real environments. In the hypothetical environment scenario, the vital rates differ between treatments, but environments (*f* and *ρ* in eqs. 4 & 5) are identical. In the real environment, both vital rates and environment differ between treatments. As (H1) examines the differences between *log* λ_s_ for each treatment at varying frequencies and autocorrelation, we tested (H1) in the hypothetical environments, constructed using a range of *f =(*0.01,…,0.99) and, *ρ* (0,…,1). Finally, to test the (H2) contribution of differences in environment and vital rates at each environment to the difference in *log* λ_*s*_ between treatments, we performed sLTRE in real environments with actual *f* and ρ, of treatments. We performed sLTRE in three cases: (i) case 1: creeping inflation (0-3%) and walking inflation (3-10%); (ii) case 2: walking inflation (3-10%) and galloping inflation (10-50%); and (iii) Case 3: galloping inflation (10-50%) and hyperinflation (>50%). In case 1, we included 17 countries in the analysis, and for cases 2 and 3, 10 and four countries, respectively. The countries selected for each case differ because, in them, both treatments and environmental states needed to have been observed.

## Result

To tested (H1), that human populations have lower stochastic growth rates when exposed to a higher inflation environment at all frequencies, and temporal autocorrelation. To do so, we used two approaches to compare long-term population growth rates among treatments. First, for each of our 76 examined countries, we calculated stochastic population growth rate *logλ*_*s*_ for a range of hypothetical environments, where the stationary frequency and temporal autocorrelation pattern of the treatments are equal, but their vital rates differ. Second, we calculate *logλ*_*s*_ for treatments under real environments, using their actual stationary frequency and temporal autocorrelation. We compared the stochastic growth rates for three cases: (i) creeping (0-3%) and walking inflation (3-10%), (ii) walking (10-50%) and galloping inflation (10-50%), and (iii) galloping and walking inflation (>50%), under both the environments.

In our first approach, we calculated *logλ*_*s*_ under hypothetical environments considering equal frequencies (0-1) and temporal autocorrelation (0-1), for each treatment. Supporting our hypothesis (H1), we found that, in case 1, the population under creeping inflation (0-3%, treatment1) has a higher *logλ*_*s*_ than that of walking inflation (3-10 %) across all the examined countries. For most (∼70%) countries, the *logλ*_*s*_ decreases with increasing frequency and temporal autocorrelation of said treatment. In countries like Belgium, Germany, Italy, and Netherlands, the *logλ*_*s*_ decreased for creeping inflation while it remained almost constant for walking inflation with increasing frequency and autocorrelation (Figure 2). Similarly, in case 2, walking inflation (3-10%) results in a higher growth rate than galloping inflation in all countries and at all frequencies and autocorrelation (Figure S1 in appendix). However, in this case, with increasing frequency of walking inflation, *logλ*_*s*_ increased for walking inflation in Lesotho, Mexico, as well as Trinidad and Tobago. Supporting our hypothesis (H1) in case 3 also, the *logλ*_*s*_ for galloping inflation (10-50%) is higher than hyperinflation (>50%) in all examined countries at all frequencies and autocorrelation, except in Argentina and Turkey (Figure S2). Moreover, with increasing frequency, the differences in stochastic growth rate between treatments in each case increase in most of the countries.

**Fig. 2.**
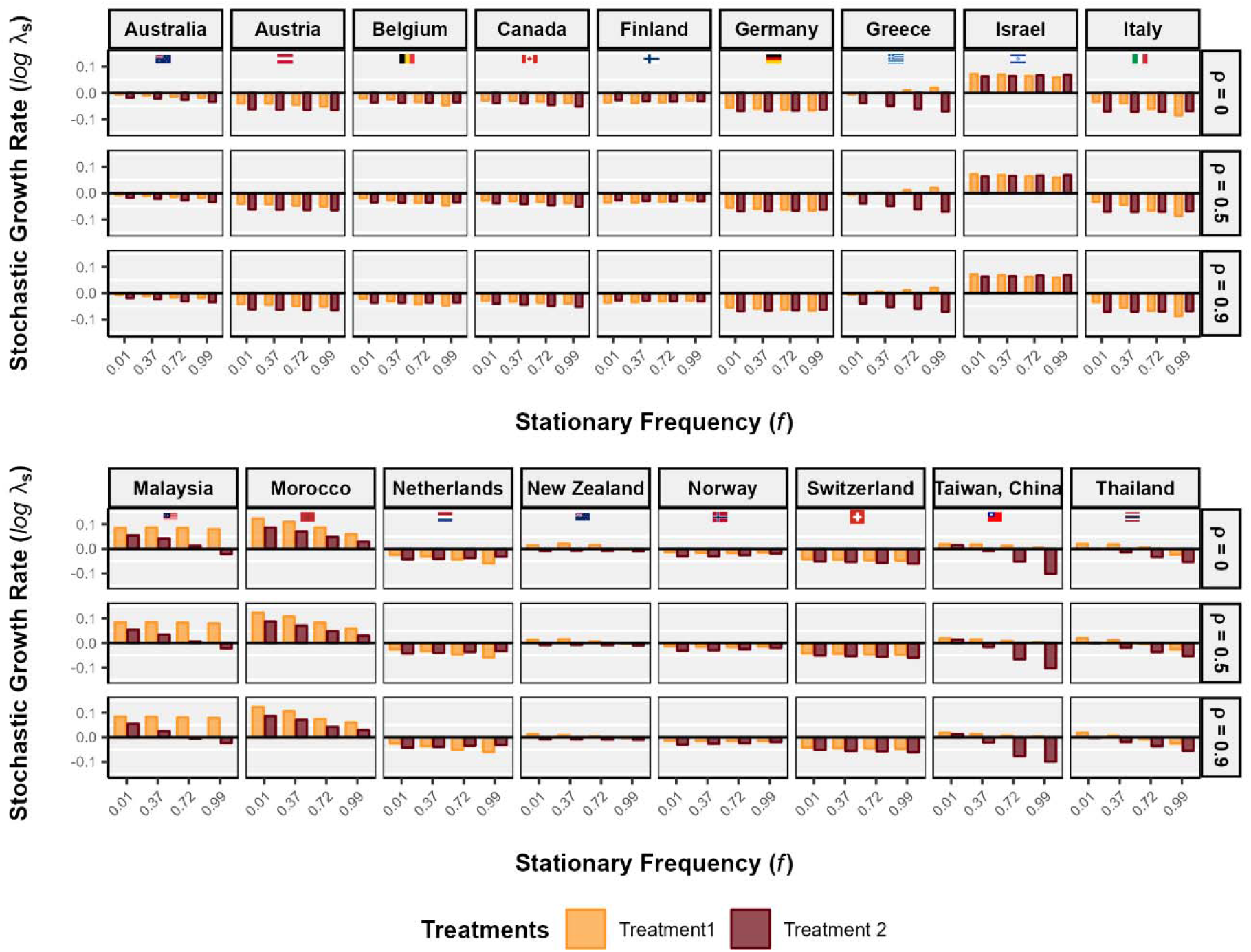
Stochastic growth rate is higher for treatment 1 (0-3%, *crippling inflation*) than that of treatment 2 (3-10%>, *walking inflation*) for all countries at all stationary frequency (*f*) and temporal autocorrelation (*ρ*) of the treatment. The stochastic growth rate decreases with increasing stationary frequency in most of the countries. However, in countries such as Germany, Italy, Netherlands, Norway, the stochastic growth rate for treatment 2 remained constant or showed small increase with increasing frequency. Here, we have presented the long-term stochastic growth rate as a function of stationary frequency (*f*) and temporal autocorrealtion (*ρ*) of the treatment. Stochastic growth rate log λ_*s*_ of treatment 1 (0-3%, *crippling inflation*) and treatment 2 (3-10%, *walking inflation*), for select countries. Growth rates are calculated by Markov chain simulation with *T* = 1000 iterations.

In our second approach, we tested whether the *logλ*_*s*_ at lower level of inflation is higher than that of higher inflation within each country at the actual frequency and autocorrelation of treatment. Supporting our hypothesis (H1), the stochastic growth rate in population at creeping inflation is systematically higher than at walking inflation in all countries examined in case 1 (Figure 3). In countries such as Greece, New Zealand, Taiwan, and Thailand, *logλ*_*s*_ at creeping inflation is positive, but it is negative at walking inflation. In case 2, *logλ*_*s*_ at walking inflation is higher than galloping inflation in all of the ten countries included in the analysis (Figure S3). In contrast to case 1, here most (∼80%) of the countries have positive *logλ*_*s*_, however, less than 0.15. Italy and Sweden have negative *logλ*_*s*_ for both treatments, whereas Chile, as well as Trinidad and Tobago have negative *logλ*_*s*_ at galloping inflation. In line with the previous cases where *logλ*_*s*_ at lower inflation has a systematic advantage over that of higher level of inflation, in case 3 also, all countries have higher *logλ*_*s*_ at galloping inflation than hyperinflation, except Argentina (Figure S4).

**Fig. 3.**
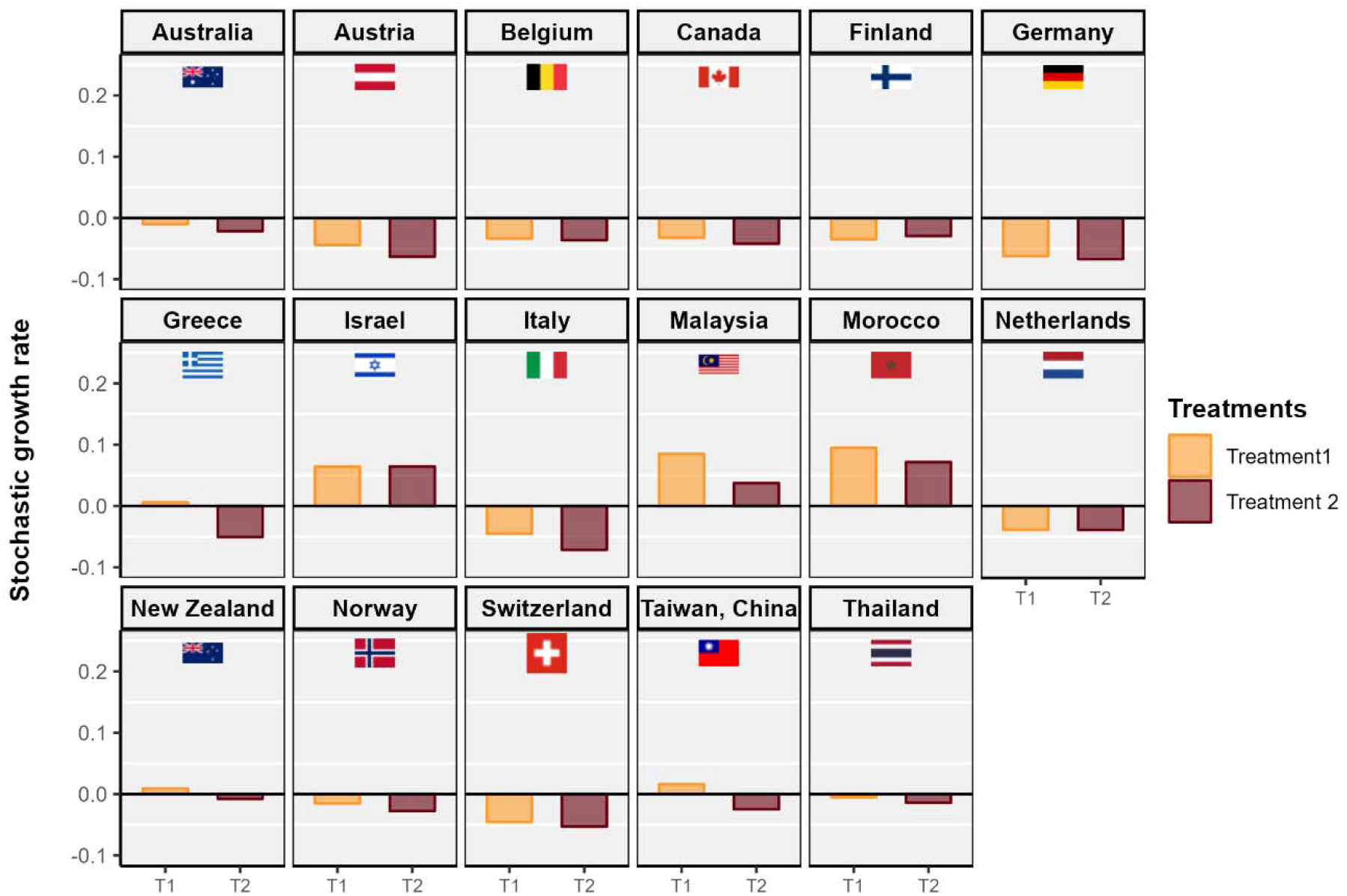
Stochastic growth rate is higher for treatment 1 (0-3%, *crippling inflation*) than that of treatment 2 (3-10%. *walking inflation*) in all countries, except in Finland. The long-term stochastic growth rate is negative in most of the countries, except in Israel, Malaysia and Morocco. Moreover, in countries such as Greece, New Zealand, and Taiwan, stochastic growth rate for treatment 1 is positive, and negative for treatment 2. The difference in stochastic growth rate between treatment 1 and 2 is high in countries like Austria, Greece, Italy, Malaysia, Morocco, Taiwan. However, the difference is very small in countries such as Belgium, Canada, Germany, Netherlands, Switzerland, and Thailand. We show the stochastic growth rate log λ_*s*_ of treatment 1 (0-3%, *crippling inflation*) and treatment 2 (3-10%, *walking inflation*), for select countries. The stochastic growth rates are calculated for variable inflation environment dictated by stationary frequency (*f*) and temporal atuocorrelation (p) of the treatments, calculated for the period 1971-2022, in the select countries. Growth rates are calculated by Markov chain simulation with *T* -1000 iterations.

To test (H2) whether lower adult survival and late age fertility at high inflation lead to lower *logλ*_*s*_, we calculated the contribution of vital rates differences to *logλ*_*s*_ differences between treatments via sLTRE. To do so, we first decomposed the differences in the *logλ*_*s*_ values across treatments in each of the three cases into contributions from matrix elements at each environmental state, stationary frequency, and temporal autocorrelation pattern of the treatments. Here also, we performed the sLTRE under two environments: (1) the hypothetical environments, in which stationary frequency (*f*, 0-1), and (2) temporal autocorrelation (*ρ*, 0-1), where *f* and ρ, are equal in each treatment. Therefore, the differences in *logλ*_*s*_ between treatments are completely determined by the differences in matrix elements (vital rates). At each frequency and temporal autocorrelation, we calculated the contribution of vital rates using environmental sensitivities of vital rates and integrated the contributions for each environmental state. Second, we have examined the contributions of the vital rates differences along with the contributions of stationary frequency and temporal autocorrelation differences between treatments to the difference in their *logλ*_*s*_ under real environments.

First, we examined the contribution of differences in vital rates within each environmental state as a function of frequency and autocorrelation of treatment under hypothetical environments. At a high frequency (*f*=1) and high temporal autocorrelation (ρ=1) the environment is overwhelmingly at state 4, and the difference in growth rate between the two treatments is contributed solely by the differences in matrices at the environmental state 4. In case 1, the contribution of matrix elements at environmental state 4 increases with increasing frequency and autocorrelation in all countries (Figure S5). At the other extreme, with *f* =0 and, ρ=1, the environment is always in state 1, and the treatment effect is due to the differences in matrix elements at state 1. In case 1, the contributions of differences in matrices at environmental states 1 has the sole contribution at *f* = 0, and moves to zero with increasing frequency. Similarly, in cases 2 and 3, with increasing frequency and autocorrelation, the contribution of differences in matrices at environmental state 4 increases rapidly while it moves to zero for that of environmental state 1 in all countries (Figure 4).

**Fig. 4.**
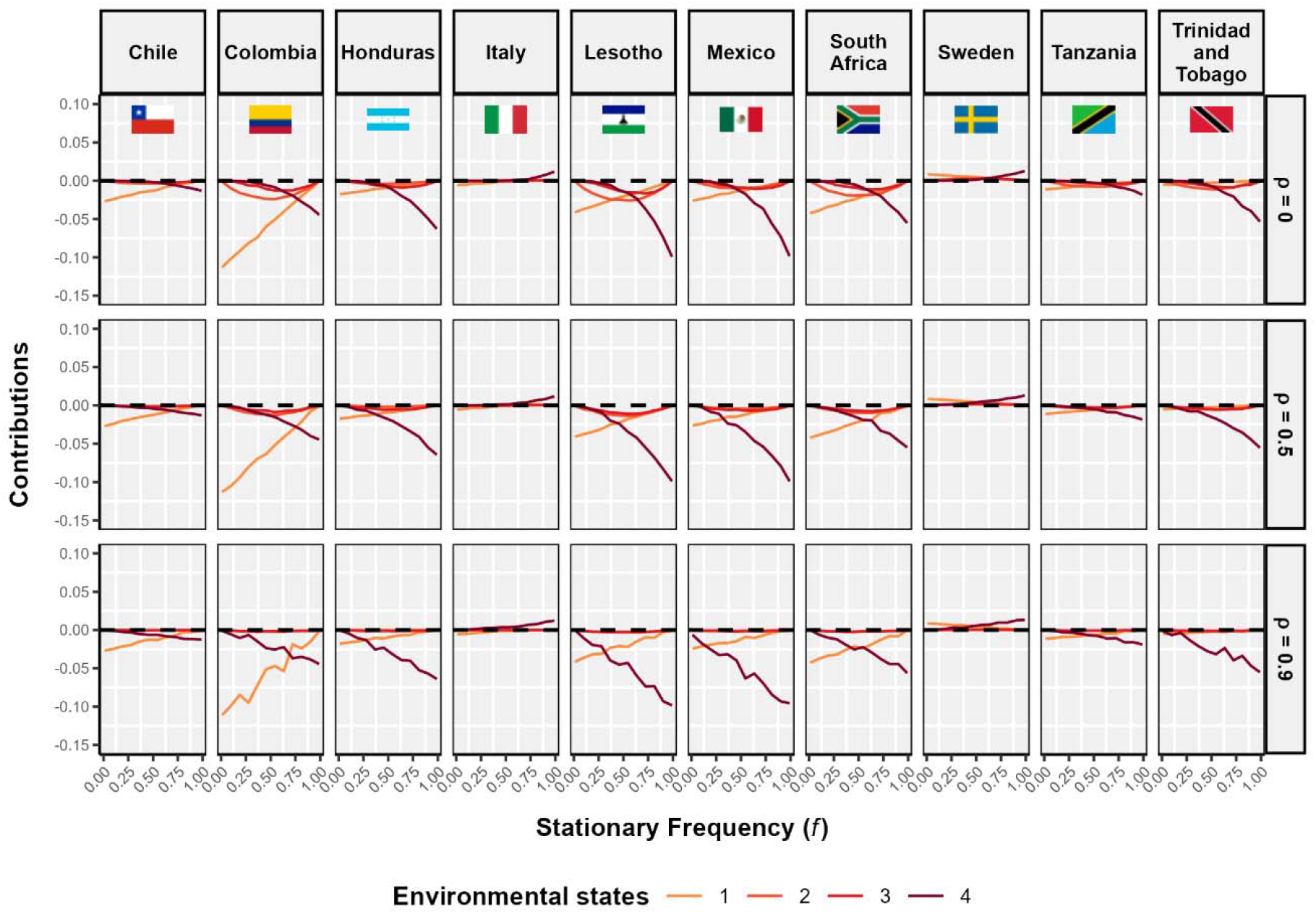
Contribution from differences in Leslie matrices at environmental state 4 **A[θ**_4_] largely determines the differences in stochastic growth rate between treatment 2 (3-10%, *walking inflation*) and treamtent 3 (10-50%, *galloping inflation*),at higher stationary frequency and temporal autocorrealtion of the treatment. With increasing frequency the contribution from **A**[**θ**_1_] moves towards zero, though with slower pace at higher temporal autocorrelation. However, the contribution from differences in matrices at environmental state 4 **A**[**θ**_4_] increases with increasing frequency, as environment is overwhelmingly at state 4 with higher frequency (*f*=0.99). we have shown the contribution of the matrices in each environmental state **A**[***θ***_*i*_] to the difference in stochastic growth rate log *λ*_*s*_ between treatment 2 (3-10%, walking inflation) and treatment 3 (10-50%, galloping inflation) as a function of stationary (*f*) and the temporal autocorrelation (ρ) of the said treatments. Growth rate and sensitivities are calculated by Markov chain simulation with T = 1000 iterations.

To examine the treatment effect, we decomposed the difference in *logλ*_*s*_ between two inflation levels into contributions from differences in frequency, autocorrelation and vital rates. The differences in vital rates at environmental state 4 had the highest contribution to the difference in *logλ*_*s*_ among all countries, except Finland, Italy, Morocco, and Thailand, in case 1 (Figure S7). In these countries differences in matrices at environmental state 1 had high contributions. The differences in frequencies and autocorrelation have a very small contribution in most of the countries, in which stationary frequency difference had a higher contribution than the former. Similarly, in case 2, the difference in vital rates at environmental state 4 between walking inflation and galloping inflation had the largest contribution to their *logλ*_*s*_ difference in all countries, except Chile, Colombia, and South Africa, where vital rates difference at environmental state 1 had the highest contributions (Figure 5). The difference in vital rates at environmental state 4 had the largest contribution to between walking inflation and galloping inflation had the largest contribution to their *logλ*_*s*_ difference between galloping inflation and hyperinflation in Argentina and Turkey (Figure S8). The vital rates at environmental state 1 contributed the most in Sudan and Venezuela.

**Fig. 5.**
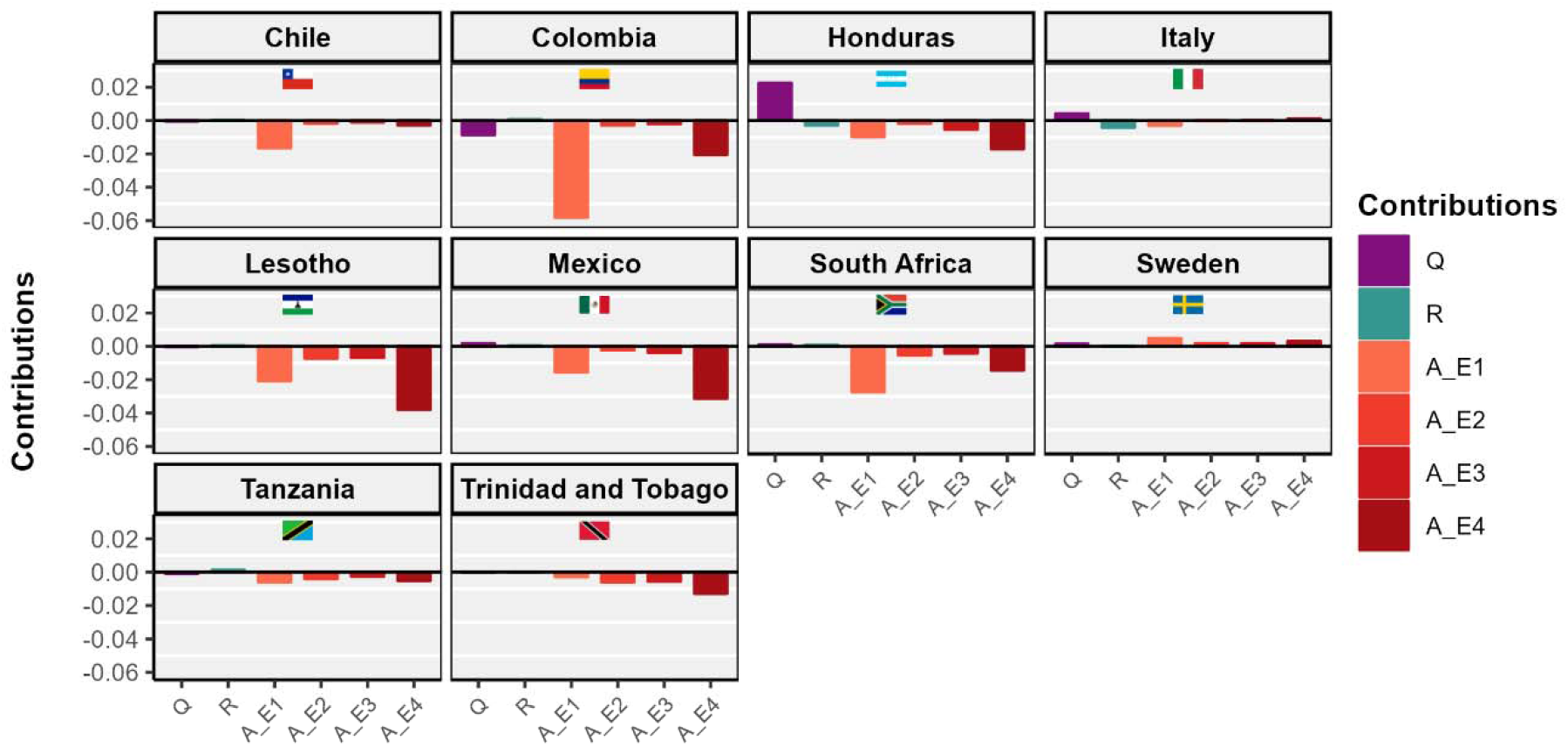
Differences in vital rates at environmental state 4 **A**[***θ***_4_] have the largest contribution to the differences in stochastic growth rate between treatment 2 {3-10%, *walking inflation*) and treatment 3 (10-50%, *galloping inflation*) for most of the countries, including Honduras, Italy, Lesotho. Mexico, Tanzania, Trinidad and Tobago. The longest brown bar shows the contribution of **A**[***θ***_4_], The contribution of the differences in matrix elements at environmental state **A**[***θ***_*j*_] to the growth rate differences is the largest in countries such as Chile, Colombia, and South Africa, shown with the largest orange bars. The contribution of environmental dynamics is very small, where long-term frequency have higher contribution than autocorrelation pattern. The purple bar shows the contribution of stationary frequency (*f*) of the treatments. We show the contributions of differences in stationary environmental frequencies (**Q**), temporal autocorrelation pattern (**R**), and vital rates (matrix elements) in each environmental state **A**[***θ***_*i*_]to differences in stochastic growth rate log A_s_ between treatment 2 (3-10%, *walking inflation*) and treatment 3 (10-50%, *galloping inflation*) Growth rates and sensitivities are calculated by Markov chain simulation with *T* = 1000 iterations.

To test whether (H2) lower-old age survival and late-age fertility determine lower at higher inflation, we identified the contribution of each matrix element differences between treatments. To do so, we have calculated the contribution of vital rates using environment-specific sensitivities via sLTRE, and summed the contribution overall environmental state. For the hypothetical environments, we have identified matrix elements which contribute to the differences in between walking inflation and galloping inflation had the largest contribution to their *logλ*_*s*_ between the two treatments in the three cases at varying stationary frequencies and temporal autocorrelation of the said treatments. Supporting our hypothesis, Figure 6 shows that in Australia, disadvantage in the survival rates at older ages, especially after 60 years and fertility rates at ages 30-45 years among the population walking inflation contributes positively to their growth disadvantage with respect to the creeping inflation. However, the contribution increases with increasing frequency and autocorrelation. Similarly, the disadvantage in survival rates at ages above 60 years at galloping inflation with respect to those in walking inflation contributes positively former’s disadvantage in long-term growth rate, while the differences in early fertility (20-25 years) contribute negatively (Figure S9). In case 3, however, the differences in fertility rates mainly contribute to the growth differences between the population at galloping inflation and hyperinflation (Figure S10).

**Fig. 6.**
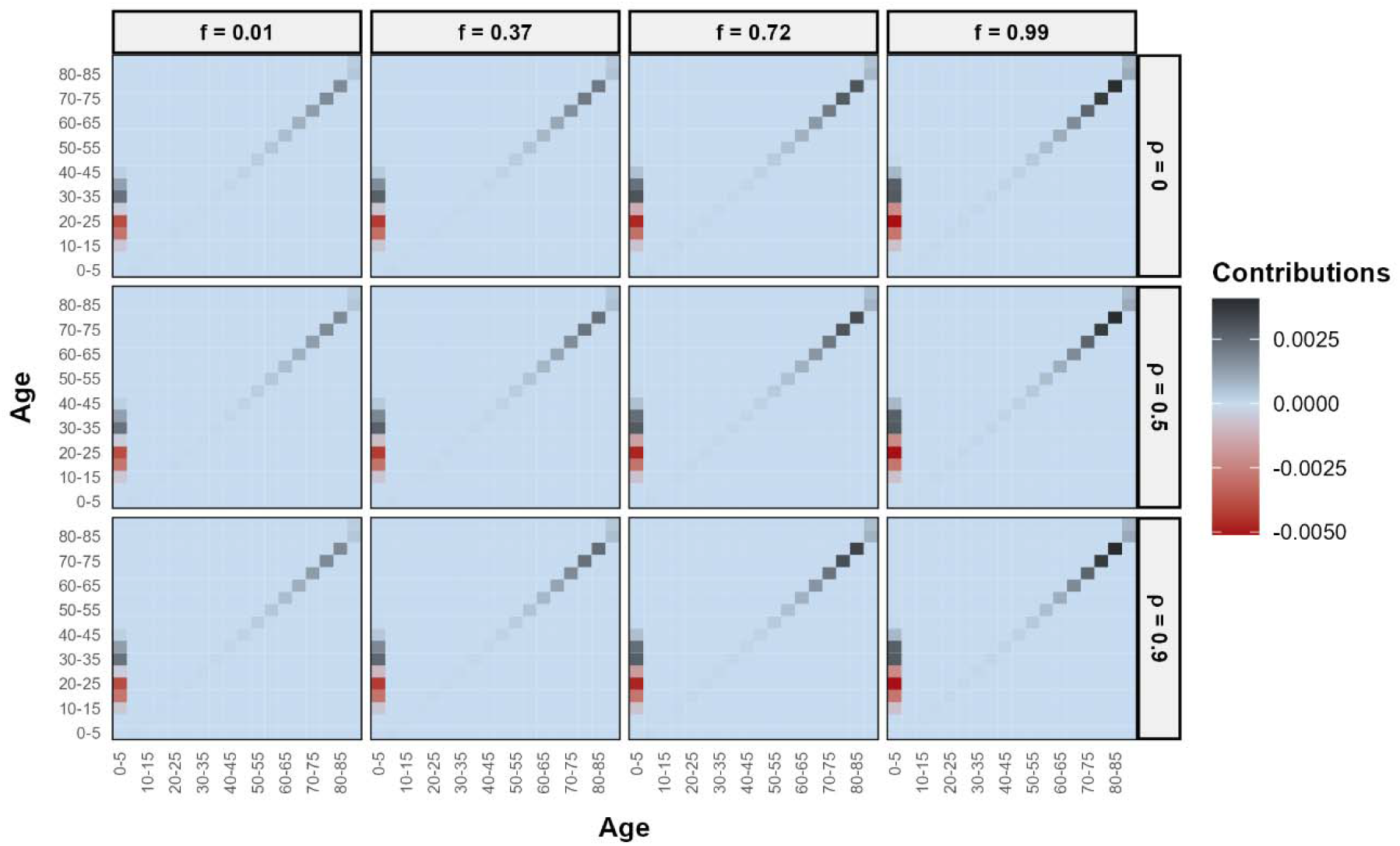
Lower survival rates at older ages (after 60 yrs) and fertility at later ages (after 30 yrs) for treatment 2 (3-10%, *walking inflation*) than treatment 1 (0-3%, *crippling inflation*) contributes positively to the differences in stochastic growth rate between treatment 1 and 2 in Australia at all stationary frequency (*f*) and temporal autocorrelation (ρ). The (positive) contribution from old age survival and late age fertility increases with increasing frequency. However, the early age fertility (15-25 years) contributes negatively to the lower stochastic growth rate for treatment 2 compared to treatment 1. Matrix plot shows the contribution of differences in matrix elements **A**_*i*_, integrated over all environmental states, to the difference in stochastic growth rate log *λ*_*s*_ of treatment 1 (0-3%, *crippling inflation*) and treatment 2 (3-10%, *walking inflation*), for Australia. Growth rate and sensitivities are calculated by Markov chain simulation with T = 1000 iterations.

For a more complete exploration of the treatment effect, we examined the contribution of vital rates in the real environment. Supporting our hypothesis (H2), lower survival rates at ages 60 years and fertility in later reproductive ages (after 30 years) at walking inflation compared to creeping inflation contributed positively to the between walking inflation and galloping inflation had the largest contribution to their *logλ*_*s*_ disadvantage in the former in all countries in case 1. However, in Morocco, juvenile survival (years 0-5) contributes positively to the between walking inflation and galloping inflation had the largest contribution to their *logλ*_*s*_ differences between creeping and walking inflation. Fertility differences in early reproductive years contributed negatively to the growth rate differences (Figure 7). In contrast, in case 2, survival disadvantage at later ages had a very small contribution to the growth disadvantage of galloping inflation over walking inflation, except in Chile, Italy and Sweden. Differences in fertility contributed negatively to the growth difference between galloping inflation and walking inflation in all countries in case 2, except Italy and Sweden (Figure S11).

**Fig. 7.**
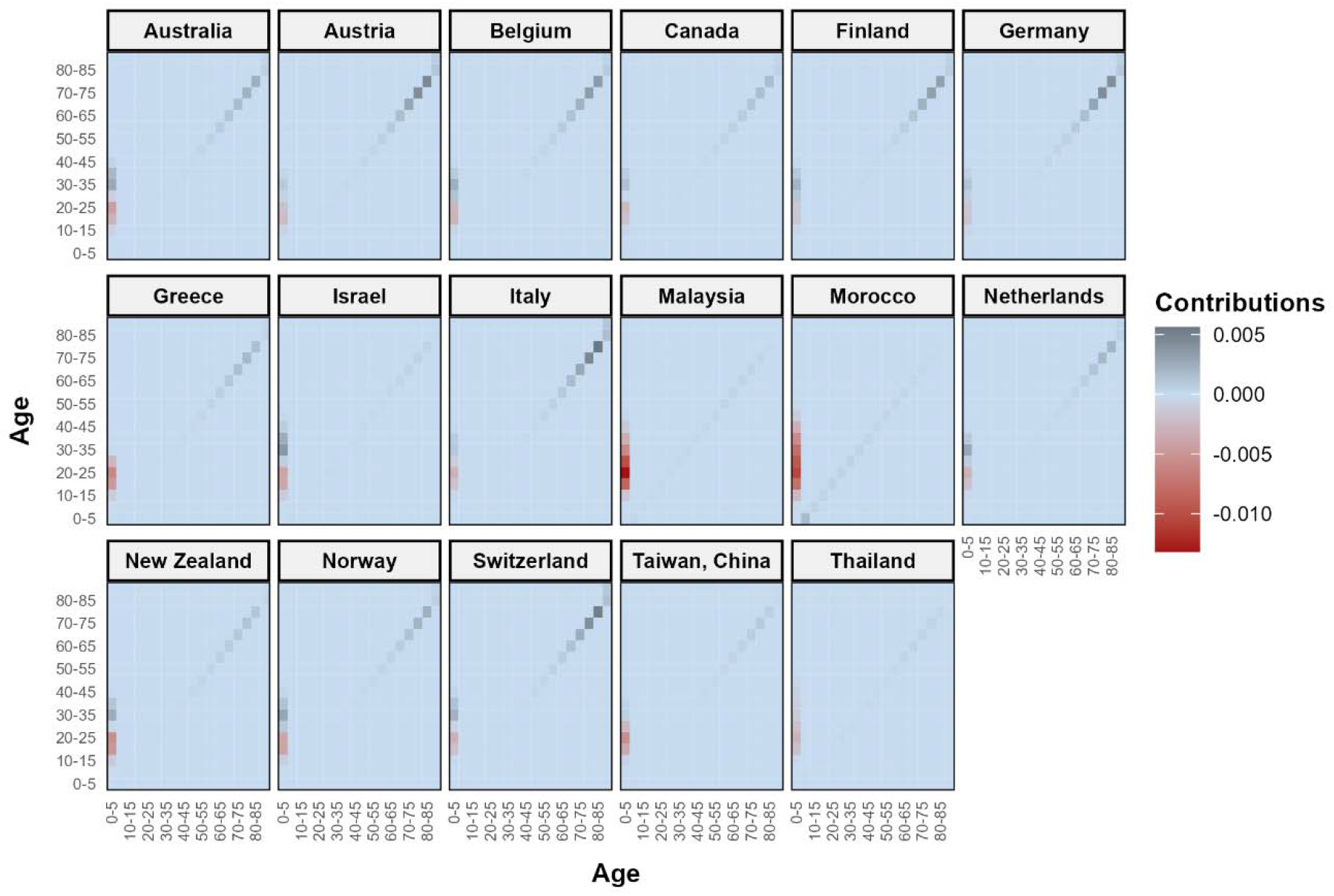
Lower adult survival rates especially at older ages (>60 yrs) and fertility at later ages (>30 yrs) for treatment 2 (3-10%>, *walking inflation*) contributes positively to the differences in stochastic growth rate disadvantage of treatment 2 over treatment 1 (0-3%, *crippling inflation*). For most of countries such as Australia, Austria, Belgium, Finland, Germany, Greece, Italy, Netherland, New Zealand, Norway, Switzerland, the contribution of adult survival increases with increasing age. The darker grey colour shows the increasing positive contribution with age in the select countries. Contrary to other countries, in Morocco higher positive contributions comes from juvenile survial. Differences in Fertility rates at early reproductive years (15-25 yrs) contributes negatively to the growth rate differences between treatment 1 and treatment 2 in all countries. The Matrix plot shows the contribution of the **A**_*i*_ integrated over all environmental states, to the difference in stochastic growth rate log *λ*_*s*_ of treatment 1 (0-3%, *crippling inflation*) and treatment 2 (3-10%, *walking inflation*), for select countries. Growth rates and sensitivities are calculated by Markov chain simulation with *T* = 1000 iterations.

Similarly, in case 3, the fertility difference had the largest and negative contribution to the difference in between walking inflation and galloping inflation had the largest contribution to their *logλ*_*s*_ between galloping and hyperinflation in all countries except Argentina (Figure S12).

## Discussion

The responses of vital rates among human populations to variability in the inflation environments determines the long-term growth rate differences between the inflation environments. Here, using stochastic growth rates *logλ*_*s*_ calculated from high-resolution inflation (Ha et al. 2023) and demographic data (United Nations, 2022) from 76 countries, we obtain support for the hypothesis that at a higher level of inflation *logλ*_*s*_ is lower than at lower inflation levels. In all three comparisons of *logλ*_*s*_ at different inflation levels, *logλ*_*s*_ is lower at a higher level of inflation in every country, except Argentina. Interestingly, with increasing frequency and temporal autocorrelation of inflation levels, the associated *logλ*_*s*_ decreases, however, more rapidly at higher levels of inflation, making the difference in *logλ*_*s*_ larger at a higher frequency. At a higher frequency of treatment, the environment is overwhelmingly at state 4, meaning the country was at the said inflation level for six or more years. As changes in vital rates among humans show lagged effects from changing inflation levels (Adsera and Menendez 2011; Juselius and Takats 2018), the differences in *logλ*_*s*_ between a low and high inflation level are more pronounced at a higher frequency and temporal autocorrelation of the treatments (*i.e*., lower variability in inflation level in the country).

Fluctuations in inflation environments are translated into population oscillations through differences in vital rates between environments. Our result highlights that differences in vital rates between treatments when the country stayed in the said inflation level for ≥6 years have the largest contributions to the differences in *logλ*_*s*_ in most of the countries. This finding signifies the fact that persistently high inflation for a duration of six years or above reduces the vital rates in a country leading to lower *logλ*_*s*_ than a lower level of inflation. The contributions from the parameter of environmental dynamics, *i.e*., stationary frequency and temporal autocorrelation of the treatments, are low in most countries, where differences in stationary frequency between treatments have a higher impact than the latter. The temporal autocorrelations cannot have an arbitrarily large effect on population dynamics, because the elements of the vital rate matrices {*A*_*i*_} must obey evolutionary and biomechanical constraints (Tuljapurkar 1982a), and for any given set {*A*_*i*_}, there are limits to population dynamics irrespective of temporal pattern (Tuljapurkar 1982a).

Humans reduce investment in adult survival, especially in old age and late-age fertility, to adapt to high inflation, thereby reducing *logλ*_*s*_ compared to low inflation.

Using sLTRE to decompose the differences in *logλ*_*s*_ between two inflation levels, we obtain support for the hypothesis that (H2) lower adult survival and fertility at late reproductive years correlates with systematically lower *logλ*_*s*_ at higher levels of inflation compared to lower levels of inflation. Furthermore, differences in fertility at early reproductive ages (15-30 years) between treatments contribute negatively to the *logλ*_*s*_ differences between them. Humans exposed to high and variable inflation levels coupled with labour market uncertainty use fertility as a life-history strategy to reduce the uncertainty in household income and, therefore, produce a higher number of children than those in low and variable inflation environments (Kohler and Kohler 2002; Sobotka et al. 2011). However, with increasing frequency, the contribution of the difference in early-age fertility decreases. We speculate that temporary unemployment shocks at low-frequency inflation reduce the opportunity cost of women’s time without affecting long-term income (substitution effect), making it a favourable condition for increasing fertility (Adsera and Menendez 2011; Butz and Ward 1979; Galor and Weil 1996; Schultz 1985; Ward and Butz 1980). Whereas, permanent unemployment associated with high-frequency persistent inflation largely reduces household permanent income (income effect), thereby negatively affecting fertility at early reproductive ages (Adsera 2005, 2011; Becker 1960, 1991; Silver 1965; Ben-Porath 1973; Yule 1906). Interestingly, in high-mortality countries such as Morocco, Colombia, South Africa, and Tanzania, differences in juvenile and child survival contribute to the differences in *logλ*_*s*_ between a high and low-inflation environment. Therefore, inflation environment fluctuations positively affect the *logλ*_*s*_ differences through the differences in adult survival and late-age fertility, and negatively through early-age fertility.

Argentina presents an exceptional case of vital rates response to variable inflation environment to the general trend observed in all other examined countries. In this country, the stochastic population growth rate, *logλ*_*s*_ at a lower level of inflation (galloping inflation, 10-50%) is lower than the *logλ*_*s*_ at a higher inflation level (hyperinflation, >50%); otherwise, in all examined countries. The possible reason behind this opposite trend is that Argentina represents a special case of inflationary periods with high inflation uncertainty (Thornton 2008). The transient period of hyperinflation can cause temporary unemployment shocks, which in turn can lead to high fertility (Adsera and Menendez 2011; Galor and Weil 1996; Schultz 1985). Because temporary unemployment shocks reduce the opportunity cost of women’s time without affecting long-term income, making it a favourable time for fertility (Adsera and Menendez 2011; Butz and Ward 1979; Galor and Weil 1996; Schultz 1985; Ward and Butz 1980). In line with this argument, our result shows that high fertility in a variable hyperinflation environment leads to an advantage in long-term stochastic growth rates in a hyperinflation environment over galloping inflation in Argentina.

Lifetime uncertainty plays a crucial role in human’s decisions to invest in survival and fertility as they have an aversion towards intertemporal inequalities in consumption. Individuals treat mortality risk as an object of intertemporal choice in a stochastic, finite lifetime (Yaari 1965) in which they determine the consumption, life-protection, and bequest plans at any age based on their survival chances at that age (Ehrlich 2000; Ehrlich and Becker 1972). Following our result, we speculate that lifetime uncertainty is high in countries at fluctuating (low frequency) high inflation levels. Therefore, it follows that humans exposed to a higher level of inflation invest less in their survival at older ages (especially above 60, where chances of survival have high uncertainty), compared to those exposed to lower inflation levels. The demand for risk aversion in mortality at a high and variable inflation environment depends on uncertainty about the return from investment in survival, discounting of present for the future utility of life years (Edwards 2013; Garcia-Sanchez and Pierrard 2023), ultimately reducing old-age survival at higher inflation levels. Similarly, we also speculate that high child mortality and lifetime uncertainty at higher levels of inflation can increase fertility at early ages. Given the convex relationship between the marginal utility of surviving children and the number of survivors, low child mortality and reduced uncertainty at a lower inflation level lead to precautionary demand for children, reducing both insurance and replacement effects on fertility (Montgomery and Cohen 1998; Preston 1978; Schultz 1969; Williams 1977); the opposite phenomenon at a high inflation level (Kalemli-Ozcan 2003, 2008; LeGrand and Sandberg 2006; Yakita 2001). Therefore, high lifetime uncertainty at a high level of inflation affects the investment in survival and fertility, subject to humans’ preference for smoothening consumption over a lifetime.

Historically, demographers have overlooked the randomness of vital rates in a variable economic environment in analysing the economic impact of population dynamics. To understand the impact of economic factors such as income, business cycle or economic growth on fertility (Adsera 2005, 2011; Bollen et al. 2007; 1960, 1991; Freedman and Thornton 1982; Simon 1969) or mortality (Deaton 2003; Rodgers 1979; Snyder and Evans 2006; Wilkinson 2022; Wilkinson and Picket 2008), researchers have considered a deterministic economic environment. However, we need to consider the response of vital rates to the fluctuations in the economic environment. Literature suggests the trend value of inflation as a variable which captures the variability in the economic environment (Ascari and Sbordone 2014; Stock and Watson 2007, 2016), which can be used to study the impact of variable economic environment on asymptotic population dynamics. Recently, attention has been paid to the impact of inflation on fertility (Chang et al. 2013; He 2018; Sobotka et al. 2011), although the impact on mortality and population growth remains almost unattained. Moreover, it is very difficult to establish the link between inflation and demography and isolate the impact of inflation on demography (Bobeica et al. 2017; Juselius and Takats 2015, 2016, 2018).

Using our framework, researchers can examine the impact of variable inflation environment on long-term population dynamics through differences in vital rates at each environment. Our framework can explicitly account for the lag effect of inflation on demographic behaviour. It also isolates the effect of inflation from the effect of other secular changes (*e.g*., technology, culture) by incorporating time-varying dynamics of the environment. Novel to this approach, we consider the long-term change in population structure (Leslie 1945), which can affect the asymptotic population dynamics in an important way. Our framework can be applied to a range of environmental drivers, such as food availability, nutrition, and cultural factors, to study the impact of variable environments on human population dynamics. The result showing the translation of vital rates response to inflation variability into long-term population growth can have profound implications for macroeconomy, long-term economic growth, human capital investment, public and private transfers in the context of changing population structure, and old-age dependency. Furthermore, our approach can be extended to a non-stationary inflation environment by taking inspiration from the transient LTRE approach developed by Koons et al. (2016). Therefore, the approach to examine the impact of variability in inflation on long-term stochastic population growth offers an improved understanding of human population dynamics in a time-varying inflation environment.

Several limitations of our framework make us cautious about the interpretation of the results. First, we use a subset of 76 countries of the world for the empirical analysis, which can mask some of the peculiar demographic responses to the variability inflation environment. However, the subset of the countries used in the analyses represents countries with low to high income and inflation levels, as well as low to high variability in inflation, which makes the result robust to most of the variable inflation environment. Second, the stochastic life table response experiment (sLTRE; Caswell 2010, 2019) approach presented here is an approximation, as it neglects well-known non-linear feedback mechanism between variability in survival, fertility, population growth and inflation environment (Caswell 2001, 2010, 2019). Third, we analysed population dynamics in a closed system, which does not consider the effect of migration on long-term population growth in variable inflation environments. However, migration can impact the population dynamics through selection, *i.e*., the fittest can leave, especially in the context of hyperinflation in Latin American countries (Rocha et al. 2021). Finally, our study attempts to isolate the impact of inflation from other secular trends by accommodating the time-varying dynamics of vital rates, across a long time-span of significant demographic, cultural and technological changes across the globe, which may limit the generalisability of the result. Therefore, the result should the interpreted cautiously by considering these limitations.

In conclusion, variability in the inflation environment affects population fluctuation through differences in vital rates between environments, more so with increasing inflation levels. Researchers examining the impact of income or inflation on vital rates using cross-sectional and small time-series data (Adsera 2005, 2011; Chang et al. 2013; He 2018; Sobotka et al. 2011) in a deterministic economic environment (Adsera 2005, 2011; Bollen et al. 2007; Deaton 2003; Freedman and Thornton 1982; Snyder and Evans 2006; Wilkinson 2022; Wilkinson and Picket 2008) are unable to illustrate the response of vital rates to variability in inflation. However, the link between inflation and demography has time-varying dynamics (Ascari and Sbordone 2014; Juselius and Takats 2015, 2016, 2018; Stock and Watson 2007, 2016), which in turn affects the long-term dynamics in human populations. Our framework exploring the impact of variable inflation environment on stochastic population growth takes into account time-varying dynamics of inflation (via stationary frequency and temporal autocorrelation pattern of inflation) and vital rates, thereby isolating the impact of inflation on population growth.

## Appendix

**Table S1.**
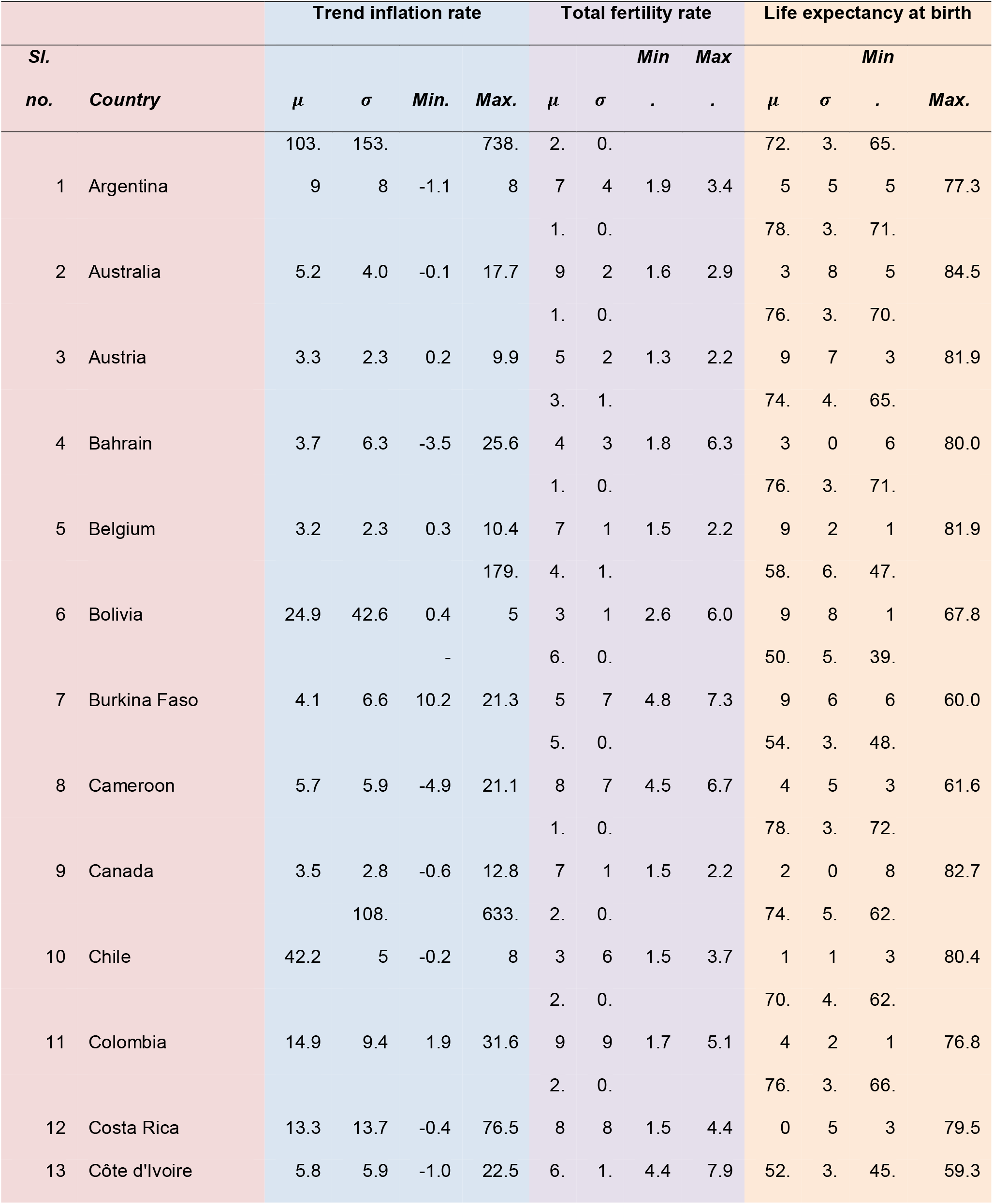
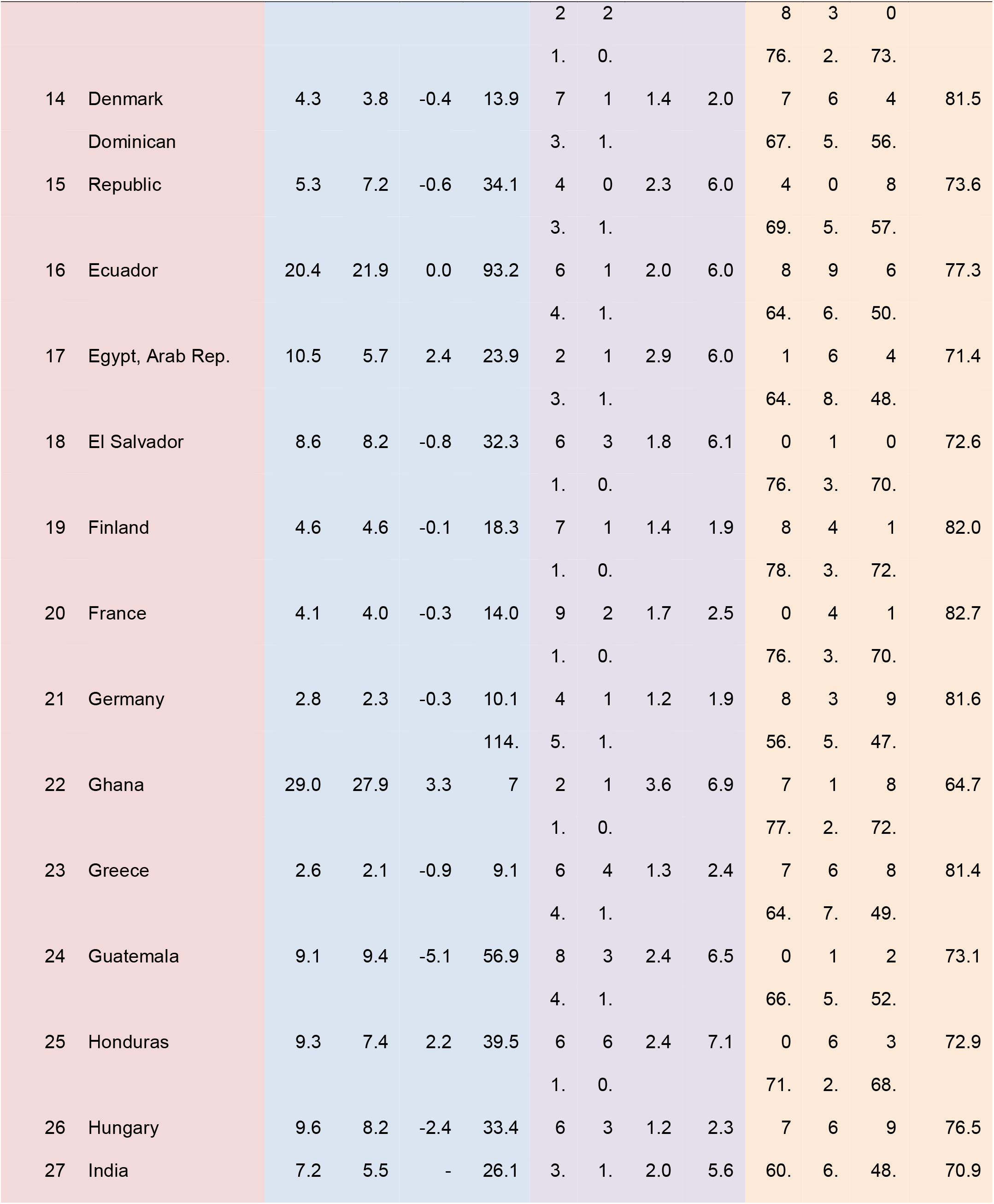
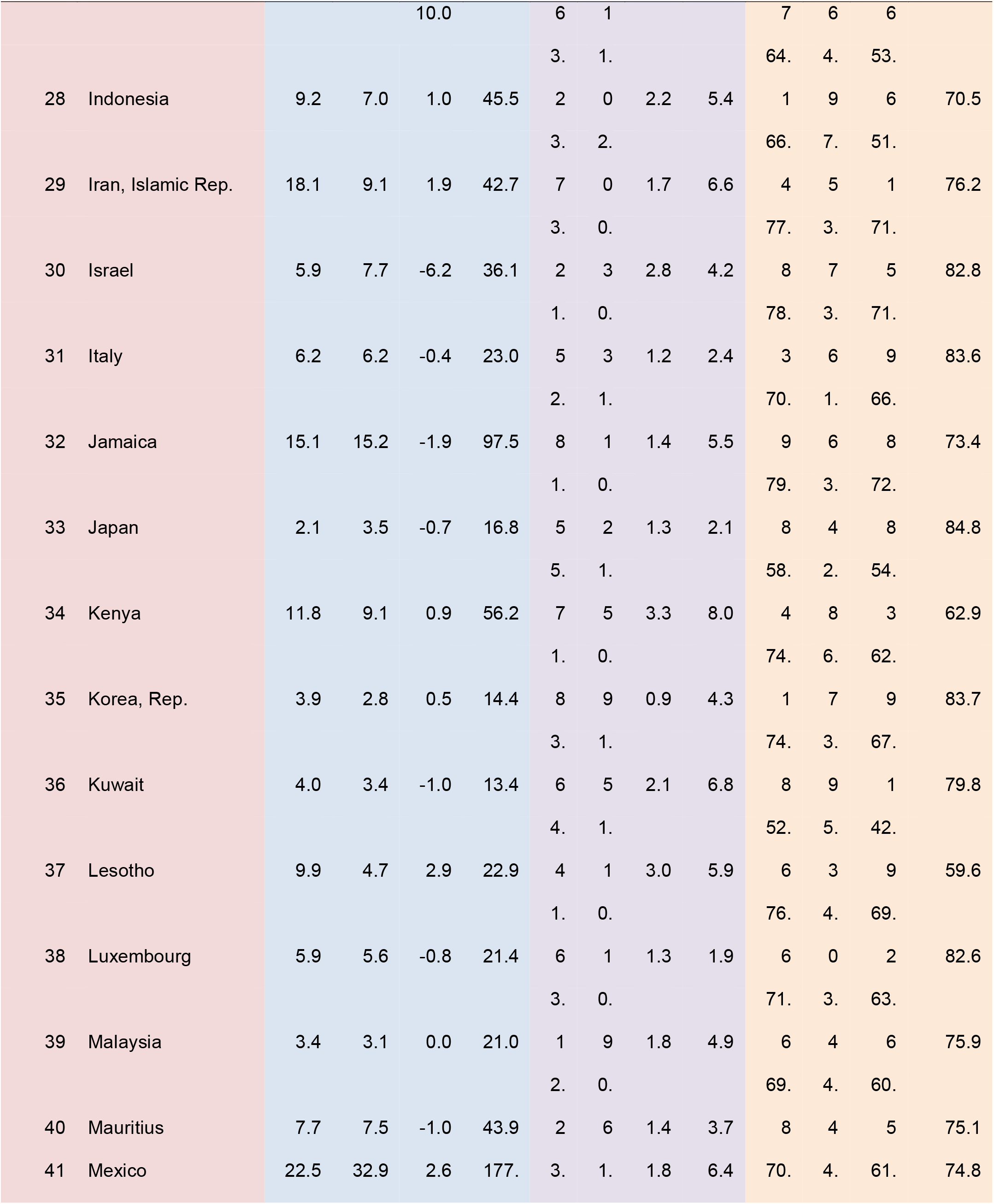
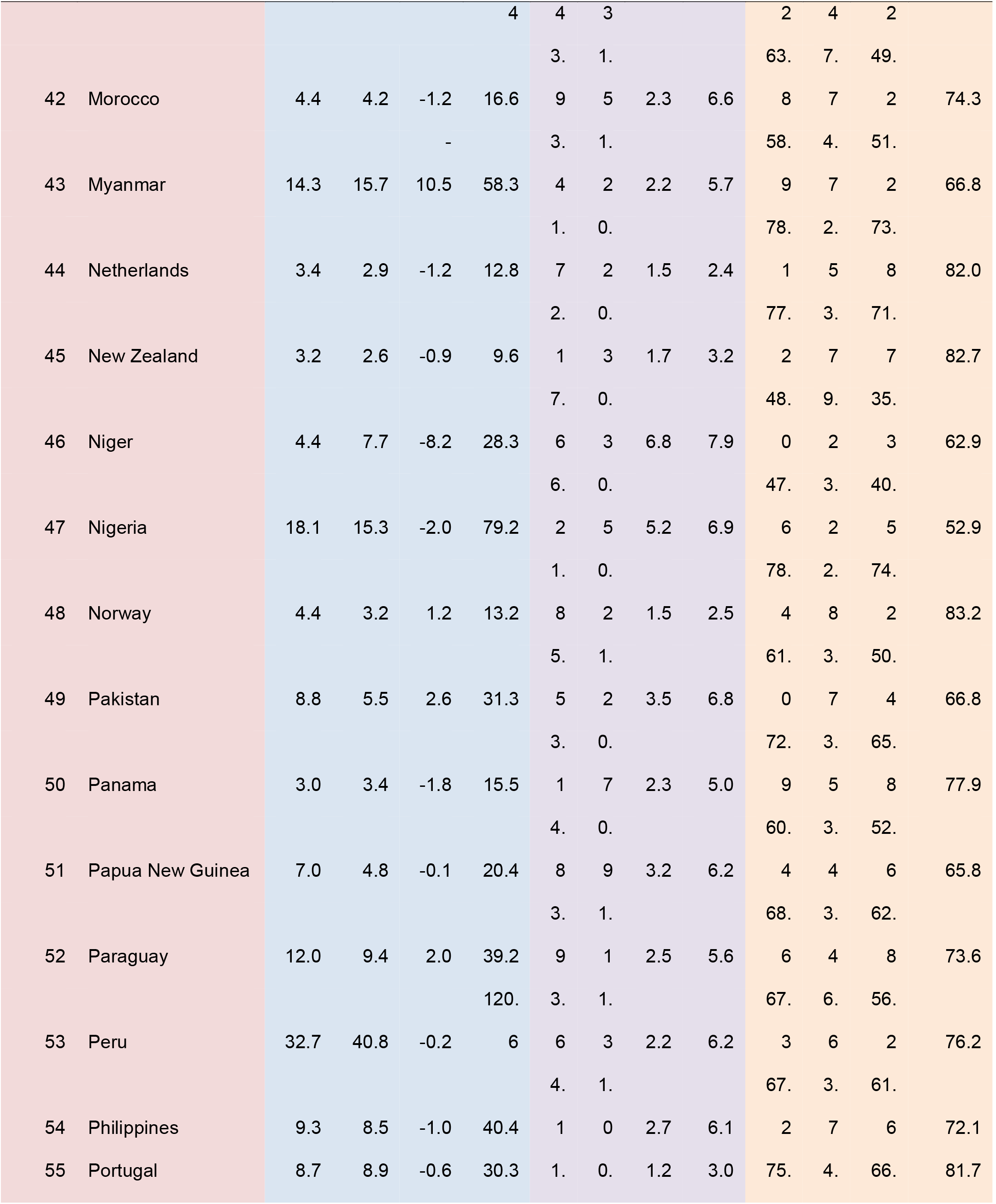
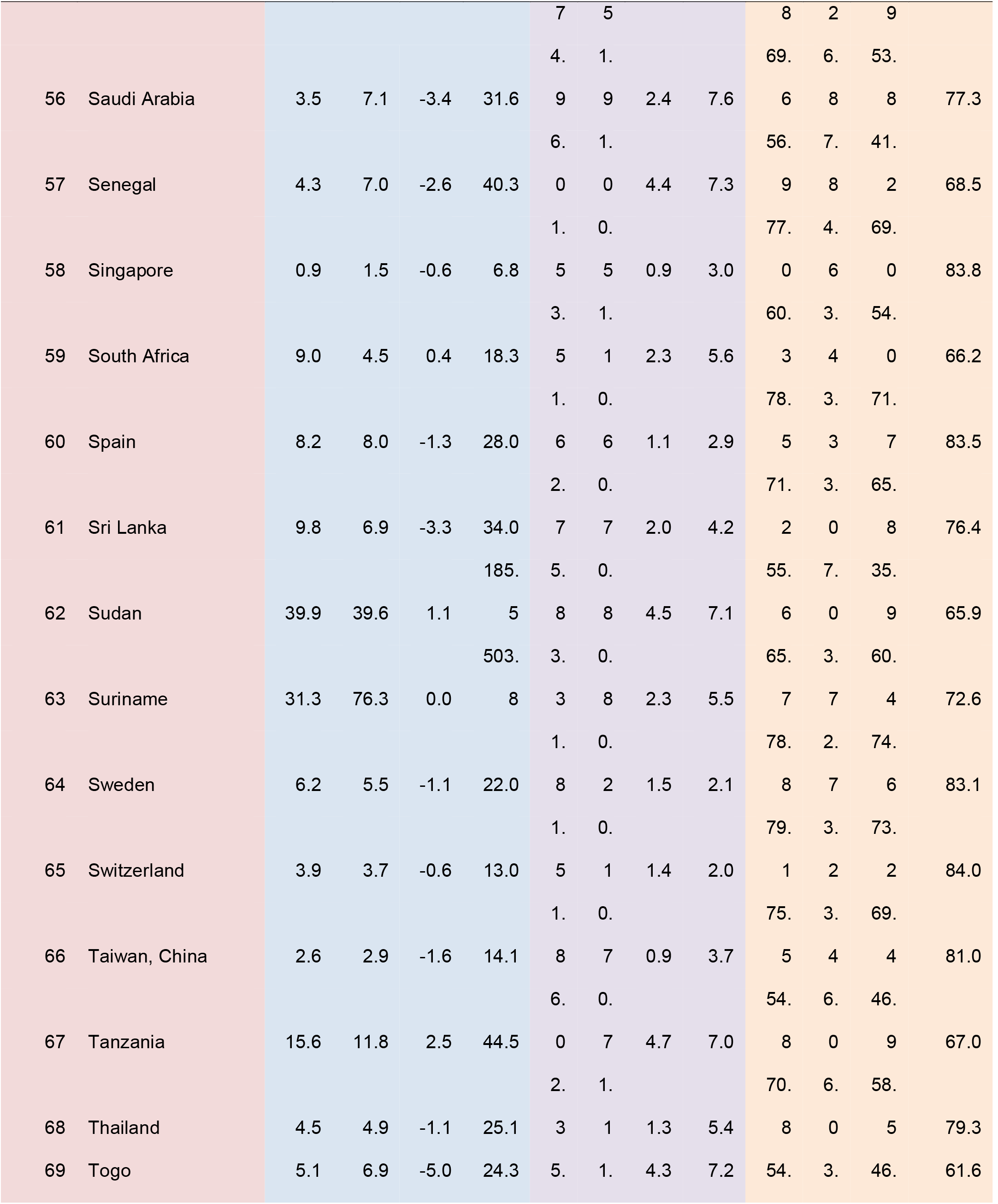
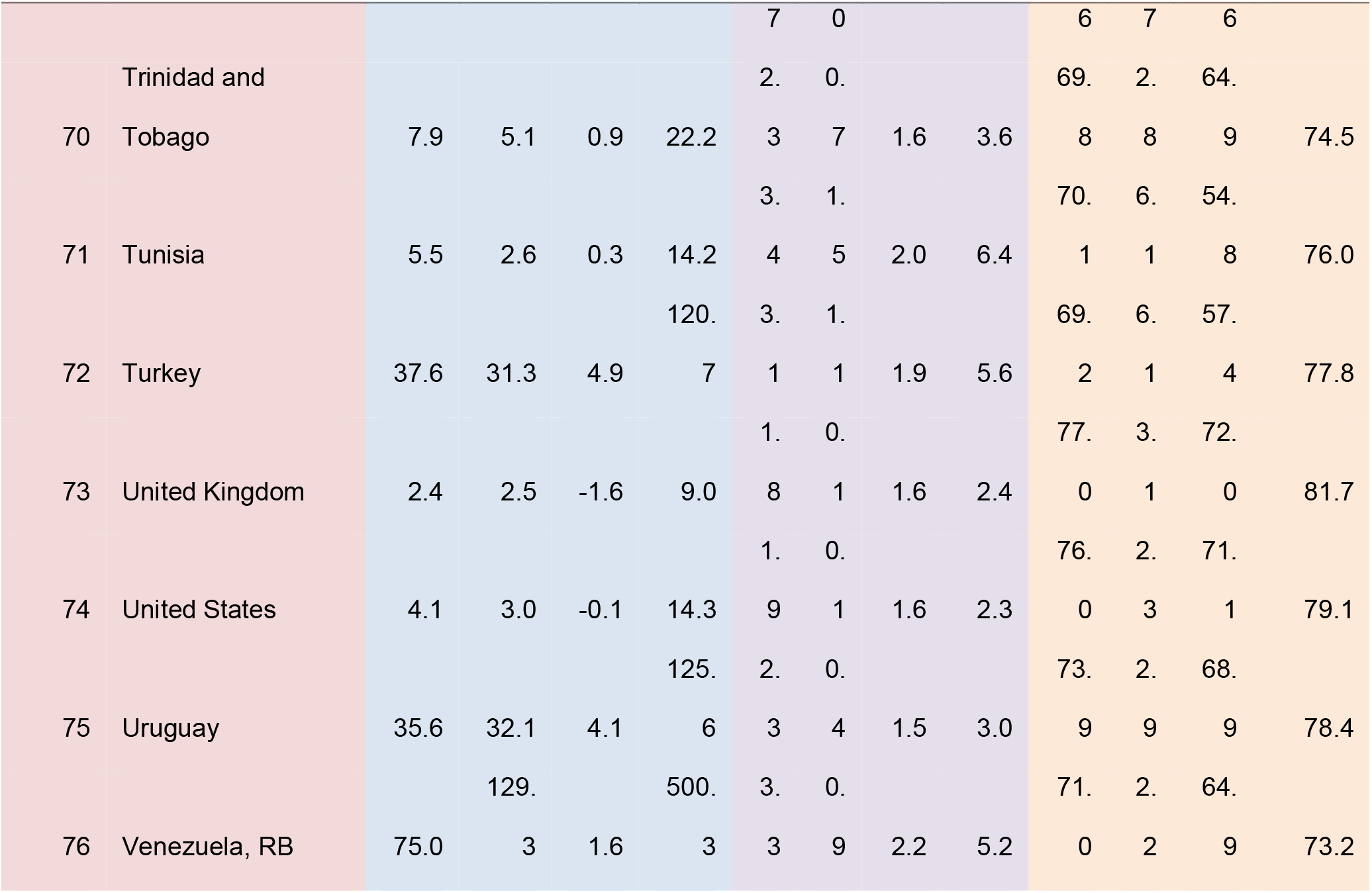
Descriptive statistics of estimated trend component of headline Consumer Price Index inflation seasonally adjusted and annualised (Ha et al. 2023), total fertility rates, and life expectancy at birth for the 76 selected countries from 1971-2022. Mean is denoted as *μ* whilst standard deviation is denoted as *σ*.

**Fig. S1.**
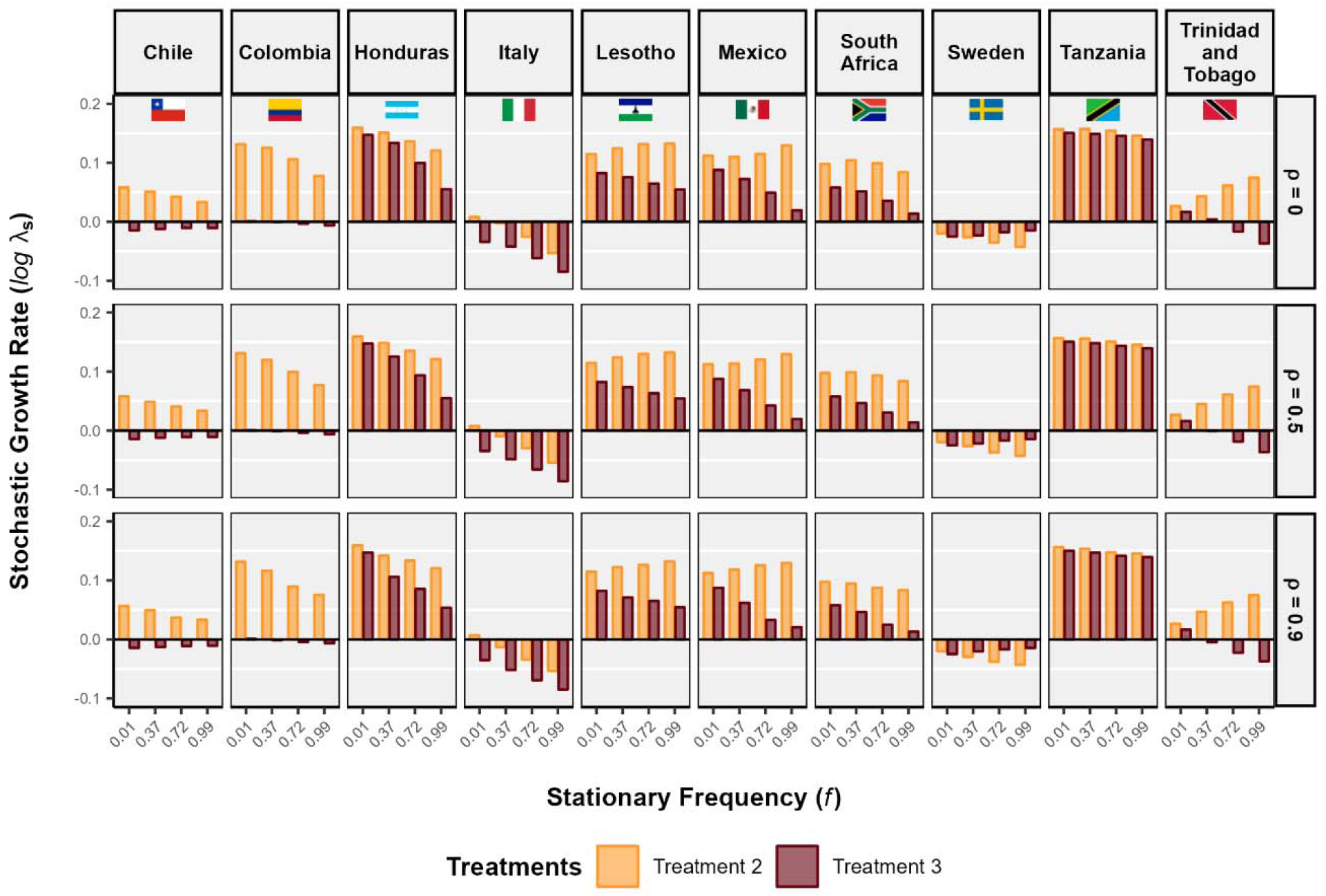
Treatment 2 *(3-10%, walking inflation)* has a growth advatange over treatment 3 *(10-50%, galloping inflation)* in all countries at all frequency (*f*) and autocorrelation (*ρ*) of treatment. The stochastic growth rate decreased with increasing stationary frequency of treatment in most of the countries. However, the difference in growth rate between treatment 2 and treatment 3 increased with increasing frequency. Countries such as Lesotho, Mexico, Trinidad and Tobago showed a different trend, where growth rate for treatment 2 increased with increasing frequency. While in Chile and Sweden growth rate for treatment 3 increased with increasing frequency. We have presented the stochastic growht rate as function of stationary frequency (*ρ*) and temporal autocorrelation λ_s_ of environment.Stochastic growth rate log A5 of treatment 2 *(3-10%, walking inflation)* and treatment 3 *(10-50%, galloping inflation)*, for select countries are calculated by Markov chain simulation with *T* = 1000 iterations.

**Fig. S2.**
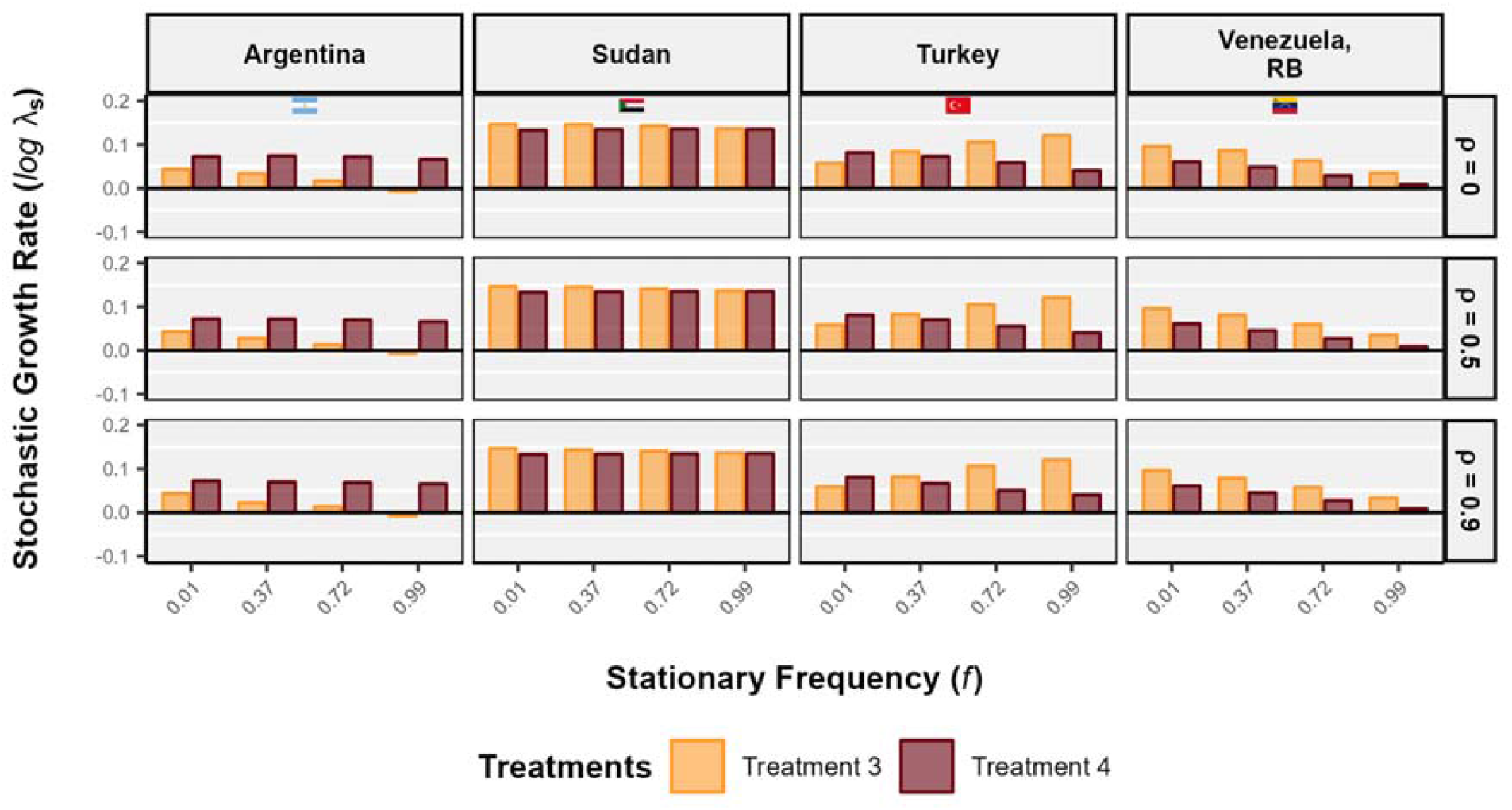
Stochastic growth rate is higher for treatment 3 (*10-50%, galloping inflation*) than that of treatment 4 (>*50%, hyperinflation*) in all countries, except in Argentina, at all stationary frequency (*f*) and temporal autocorrelation (*ρ*) of treatments. With increasing frequency, stochastic growth rate decreases for most of the countries. However, in Turkey the stochastic growth rate for treatment 3 increased with increasing frequency, making the stochastic growth rate for treatment 3 higher than treatment 4 at higher frequencies. In Argentina and Sudan,the stochastic growth rate for treatment 4 remained almost constant, while the stochastic growth rate for treatment 3 decreased. Therefore, decreasing stochastic growth rate for treatment 3 and constant growth rate for treatment 4 makes their differece higher for Argentina and lower for Sudan, with increasing frequency. In Turkey, decreasing stochastic growth rate for treatment 4 and increasing growht rate for treatment 3 makes their difference larger with increasing frequency. Here, we present stochastic growth rate as a function of stationary frequency and temporal autocorrelation of the treatments. Stochastic growth rate log λ_s_ of treatment 3 (*10-50%, galloping inflation*) and treatment 4 (>*50%, hyperinflation*), for select countries. Growth rates are calculated by Markov chain simulation with *T* = 1000 iterations.

**Fig. S3.**
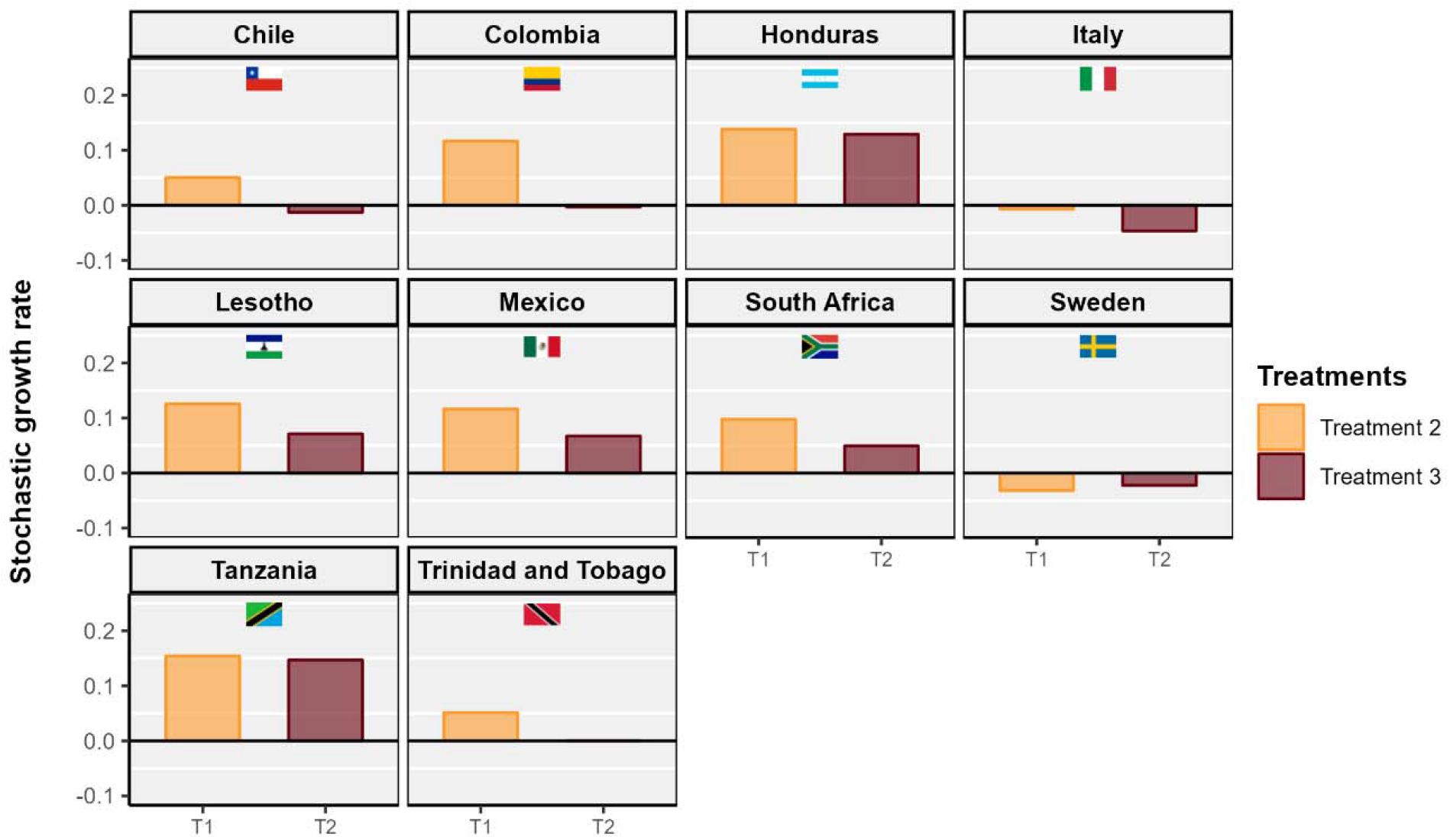
Stochastic growth rate is higher for treatment 2 *(3-10%, walking inflation)* than that of treatment 3 (*10-50%, galloping inflation)* in all countries, except Sweden. The stochastic growth rate positive for almost all countries except Sweden and Italy. Moreover, countries such as Chile Colombia, Trinidad and Tobago has negative stochastic growth rate for treatment 3. The difference in stochastic growth rate for treatment 2 and 3 is high in countries such as Chile, Colombia, Lesotho, Mexico, South Africa, Trinidad and Tobago. However, In Honduras, Italy, and Tanzania the difference is very small. We show the stochastic growth rate log λ_s_ of treatment 2 *(3-10%, walking inflation)* and treatment 3 (*10-50%, galloping inflation)*, for select countries. The stochastic growth rates are calculated for variable inflation environment dictated by stationary frequency *(f)* and temporal atuocorrelation (*ρ*) of the treatments, calculated for the period 1971-2022, in the select countries. Growth rates are calculated by Markov chain simulation with *T* = 1000 iterations.

**Fig. S4.**
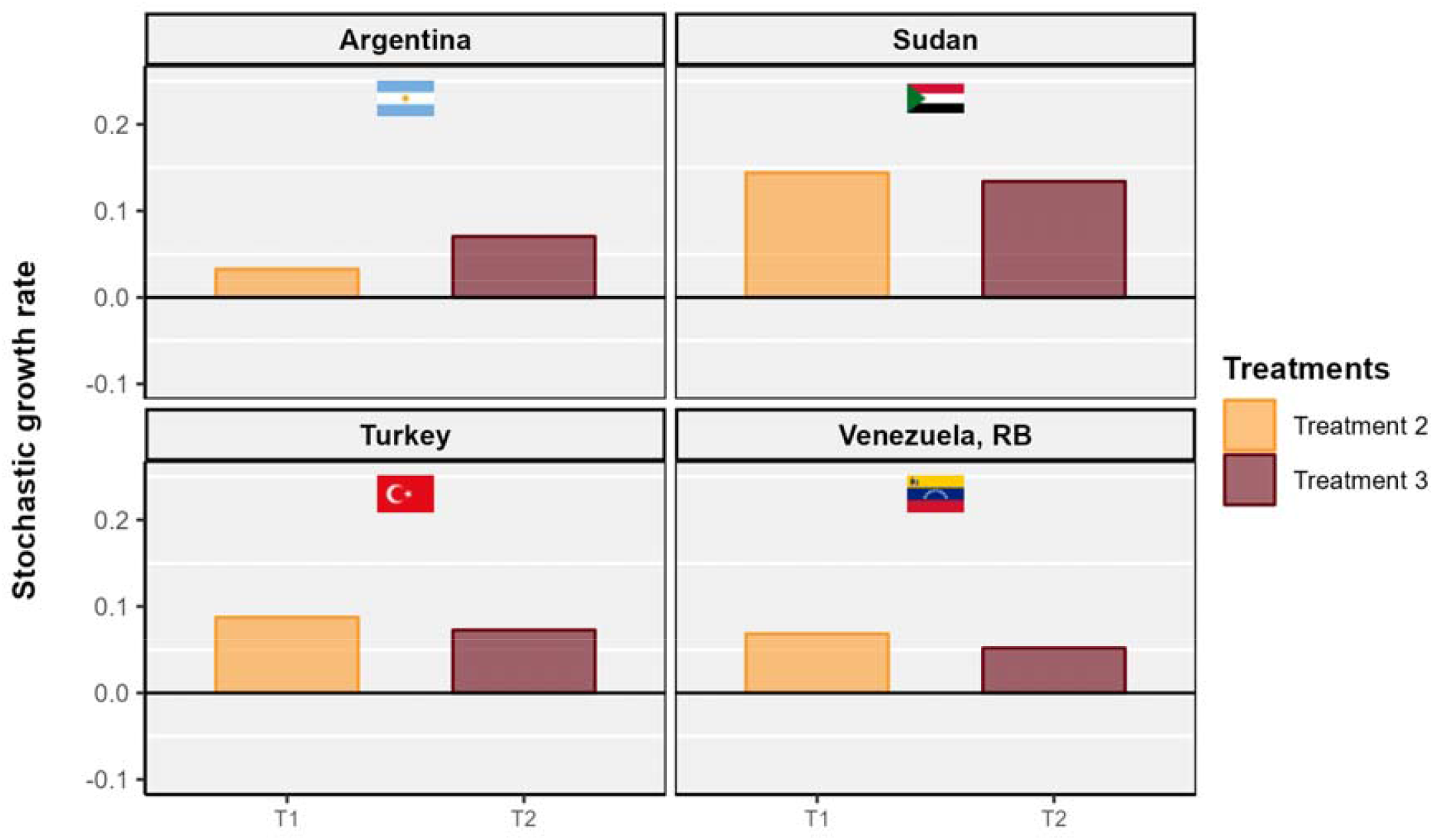
Stochastic growth rate is higher for treatment 3 (*10-50%, galloping inflation)* than that of treatment 4 *(>50%, hyperinflation)* in all countries, except in Argentina. However, the differences in the stochastic growth rate between treatment 3 and 4 is smaller than previous cases. Stochastic growth rate log λ_s_ of treatment 3 *(10-50%, galloping inflation)* and treatment 4 *(>50%, hyperinflation)*, for select countries. The stochastic growth rates are calculated for variable inflation environment dictated by stationary frequency (f) and temporal atuocorrelation (p) of the treatments, calculated for the period 1971-2022, in the select countries. Growth rates are calculated by Markov chain simulation with *T* = 1000 iterations.

**Fig. S5.**
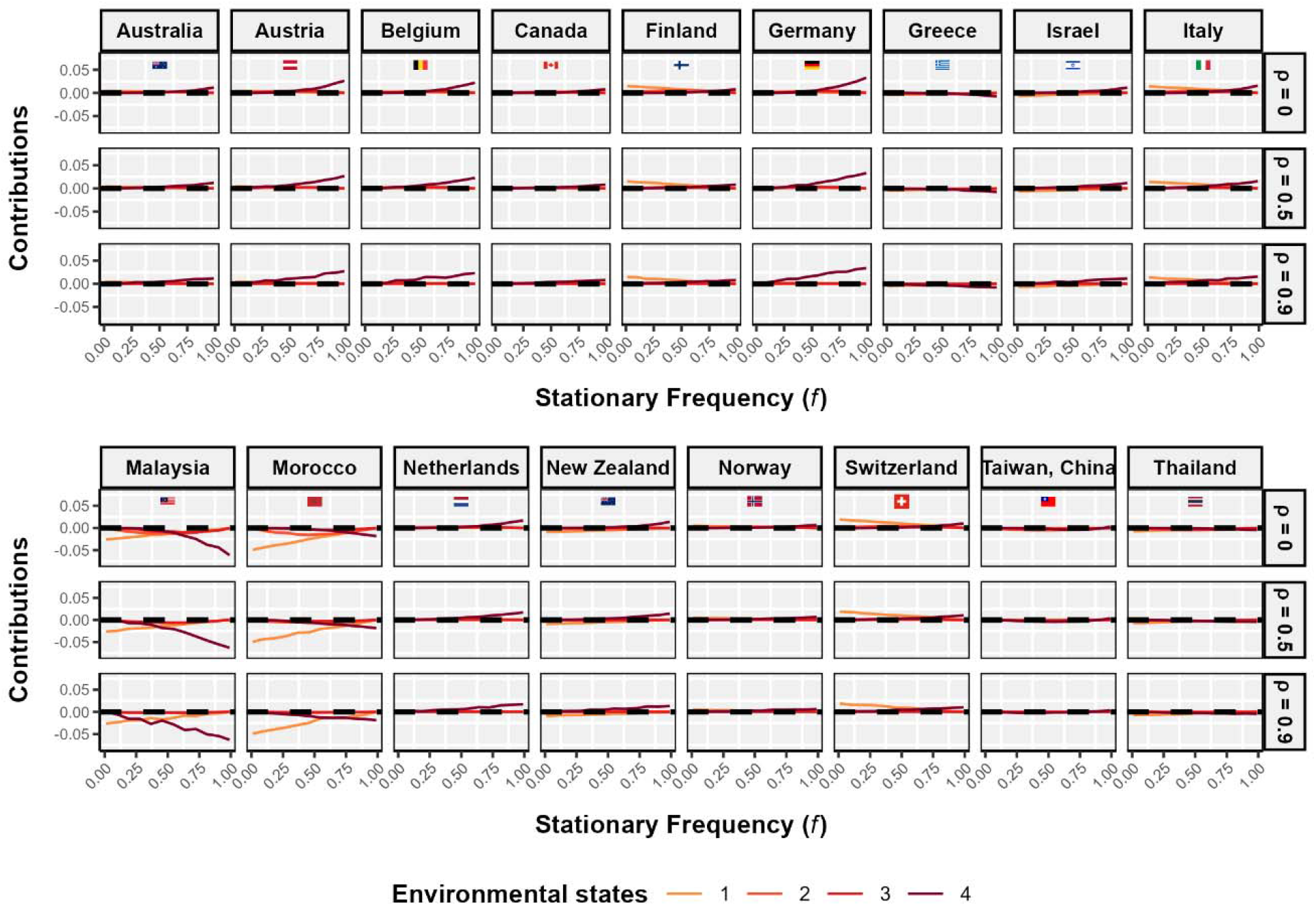
Differences between matrices at environmental states 4, **A[θ**_***4***_], contributes most to the difference in stochastic growth rate between treatment 1 *(0-3%, crippling inflation)* and treatment 2*(3-10%, walking inflation)*. With increasing stationary frequency and temporal autocorrelation of treatments, the environment is overwhelmingly at state 4. Therefore, the contribution comes from differences in A[8.J between treatments. In Countries like Malaysia Morocco, the contribution matcires at environmental state 4 **A[θ**_***4***_] increases while that in environmental state 1 **A[θ**_***1***_] decreases with increasing frequency and temporal autocorrelation. Contribution of the matrices in each environmental state **A[θ**_***i***_] to the difference in stochastic growth rate log ρ between treatment 1 *(0-3%, crippling inflation)* and treatment 2 *(3-10%, walking inflation)* as a function of stationary *(f)* and the temporal autocorrelation (*ρ*) of the said treatments. Growth rate and sensitivities are calculated by Markov chain simulation with T = 1000 iterations.

**Fig. S6.**
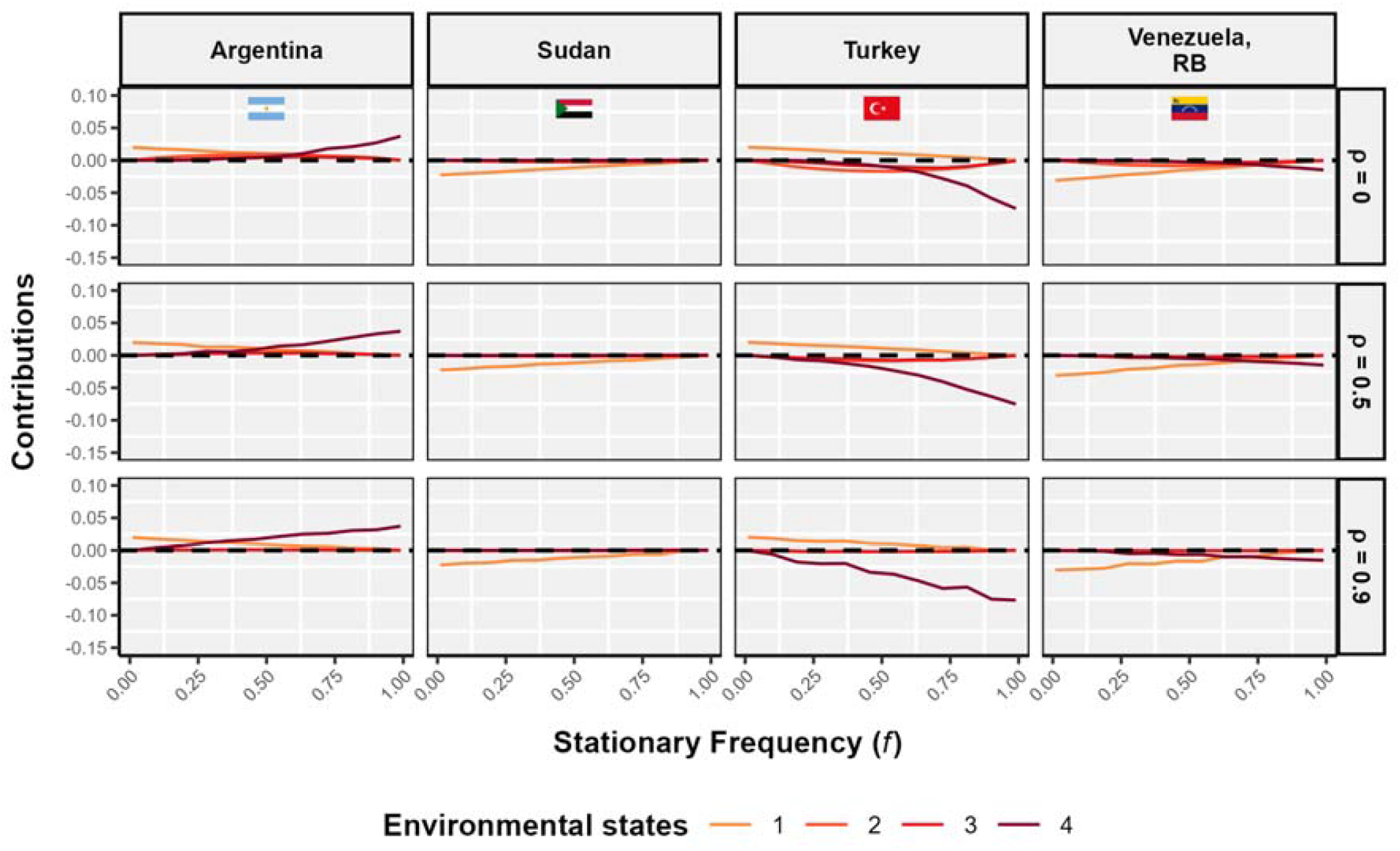
Contribution from differences in Leslie matrices in environmental state 4 **A[84)** between treatment 3 (*10-50%, galloping inflation)* and treatment 4 (>50%, *hyperinflation)* is negative in all countries, except Argentina and increases with incresing frequency and autocorrelation. At frequency *f* = *1* the contribution is completely from the environmental state 4. However, with increasing frequency the contribution from environmental state 1 **A[θ**_***1***_) decreases and is negative for all countries, except in Argentina and Turkey. At frequency *f* = *0* the environement is in state 1 and the differences in stochastic growth rate between treatment 3 and 4 completely due to differences in the matrices at environmental state 1 **A[θ**_***1***_). Here, we have presented the contribution of the matrices in each environmental state **A[θ**_***i***_) to the difference in stochastic growth rate log A5 between treatment 3 *(10-50%, galloping inflation)* and treatment 4 (>50%, *hyperinflation)* as a function of stationary *(f)* and the temporal autocorrelation (*ρ*) of the said treatments. Growth rate and sensitivities are calculated by Markov chain simulation with T = 1000 iterations.

**Fig. S7.**
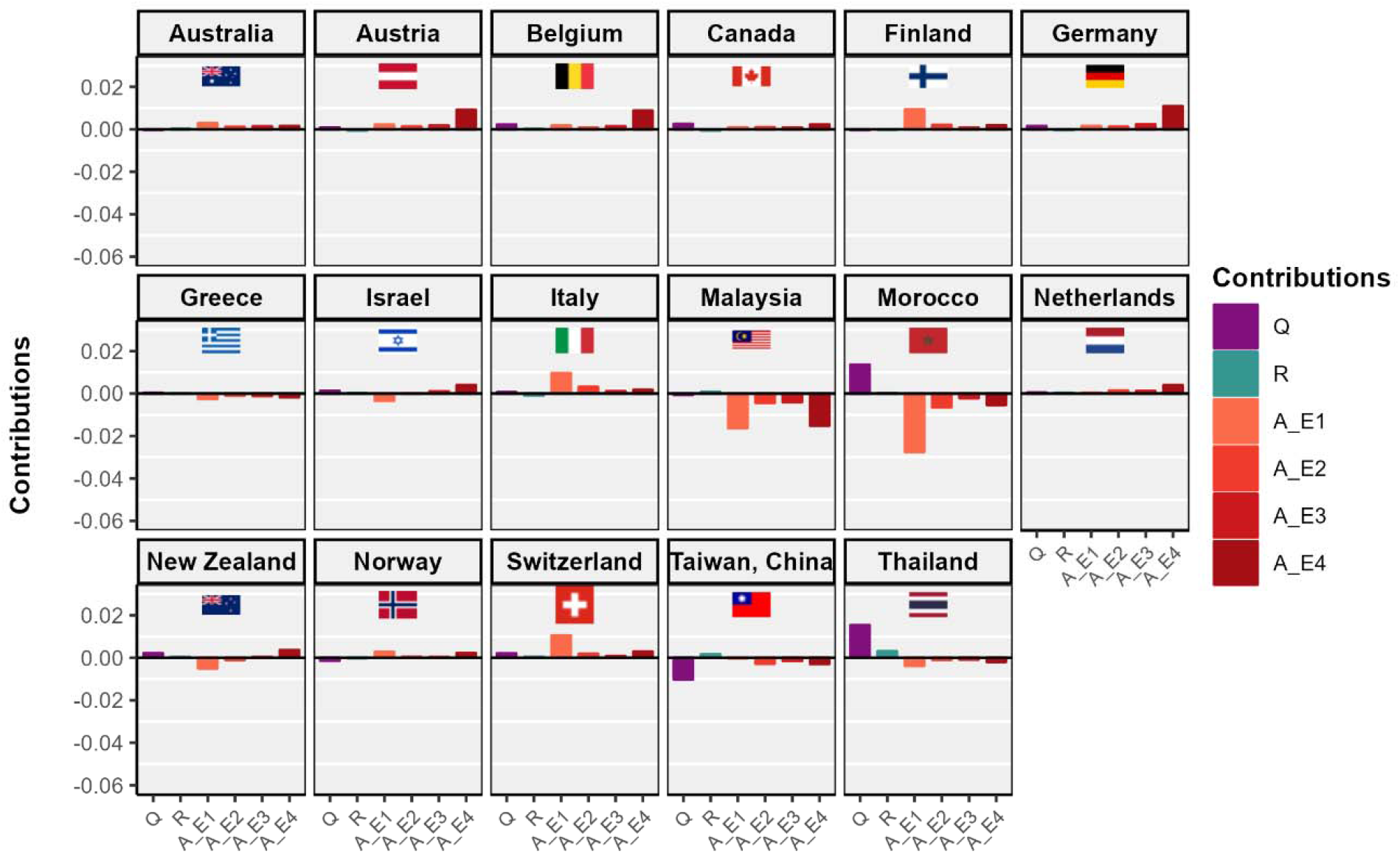
Differences in vital rates at environmental state **4 A[θ**_***4***_] have the largest contribution to the differences in stochastic growth rate between treatment 1 (0-3%, *crippling inflation)* and treatment 2 *(3-10%, walking inflation)* for most of the countries, such as Austria, Belgium, Canada, Germany, Israel, Netherlands, New Zealand. Contributions from the differences in vital rates at environmental state 1 **A[θ**_***1***_] had the largest contribution in countries including Finland, Italy, Malaysia, Morocco, Switzerland. The contribution of differences in environmental dynamics is very small, where the differences in stationary frequency (f) have higher contribution to the growth rate differences between treatment 1 and 2 than that of temporal autocorrelation pattern *(p)*. We show the contributions of differences in stationary environmental frequencies **(Q)**, temporal autocorrelation pattern **(R)**, and vita l rates (matrix elements) in each environmental state **A[θ**_***i***_] to differences in stochastic growth rate log A5 between treatment 1 (*0-3*%, *crippling inflation)* and treatment 2 *(3-10%, walking inflation)*. Growth rates and sensitivities are calculated by Markov chain simulation with *T* = 1000 iterations.

**Fig. S8.**
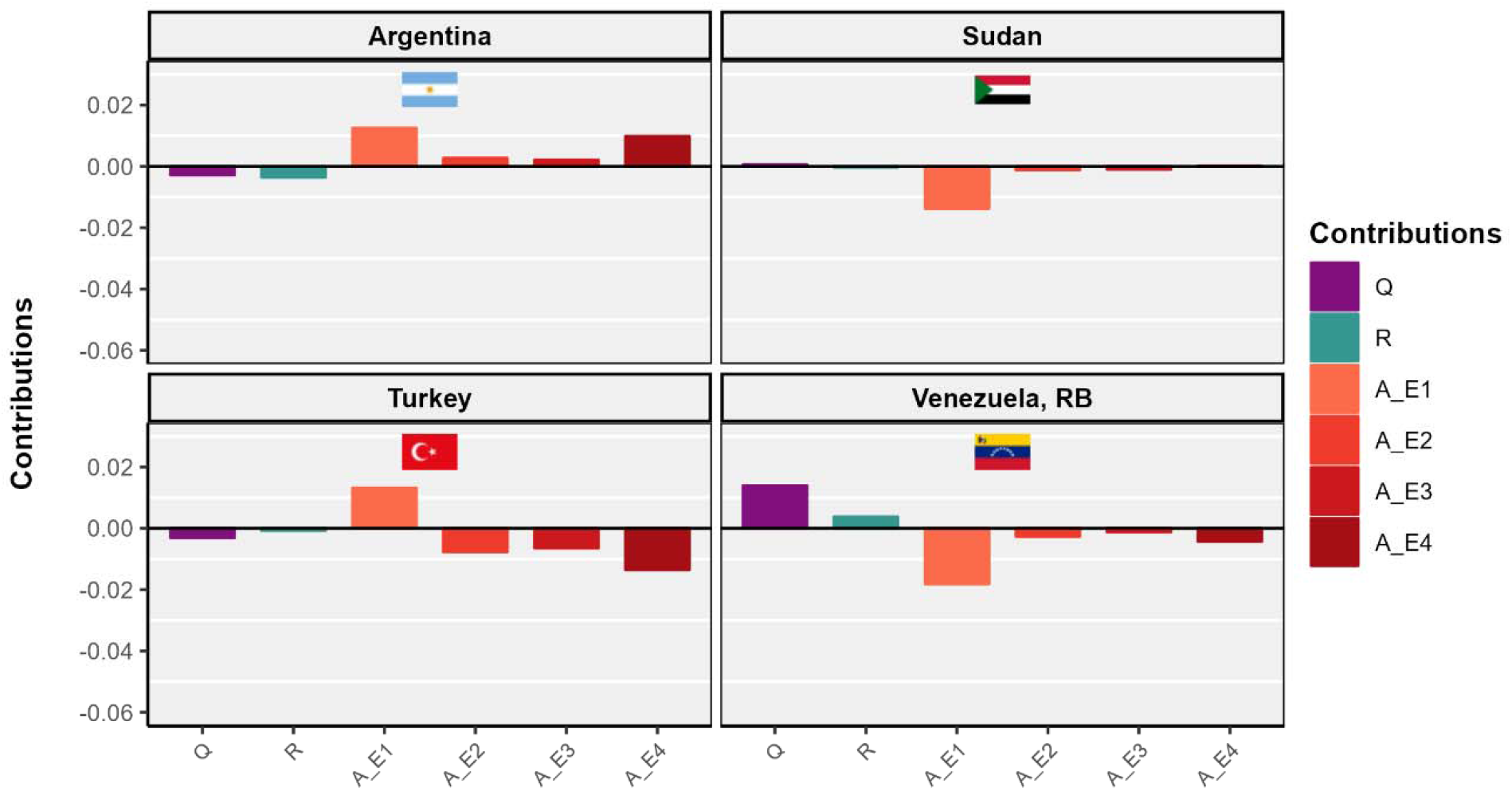
Differences in vital rates at environmental state 1 **A[θ**_***1***_] have the largest contribution to the differences in stochastic growth rate between treatment 3 *(10-50%, galloping inflation)* and treatment 4 (>50%, *hyperinflation)* for most of the countries. The contribution of matrices at environmental state 1 **A[θ**_***1***_] is shown in the orange bar. The contribution of the matrices at environmenta l state 4 **A[θ**_***4***_] is the second highest, shown in the brown bar. The contribution of environmental dynamics is very small, where long-term frequency have higher contribution than autocorrelation pattern. Contribution of the stationary frequency is shown by the purple bar. We show the contributions of differences in stationary environmental frequencies **(Q)**, temporal autocorrelation pattern **(R)**, and vital rates (matrix elements) in each environmental state **A[θ**_***i***_] to differences in stochastic growth rate log λ_s_ between treatment 3 *(10-50%, galloping inflation)* and treatment 4 (>50%, *hyperinflation)*. Growth rates and sensitivities are calculated by Markov chain simulation with *T* = 1000 iterations.

**Fig. S9.**
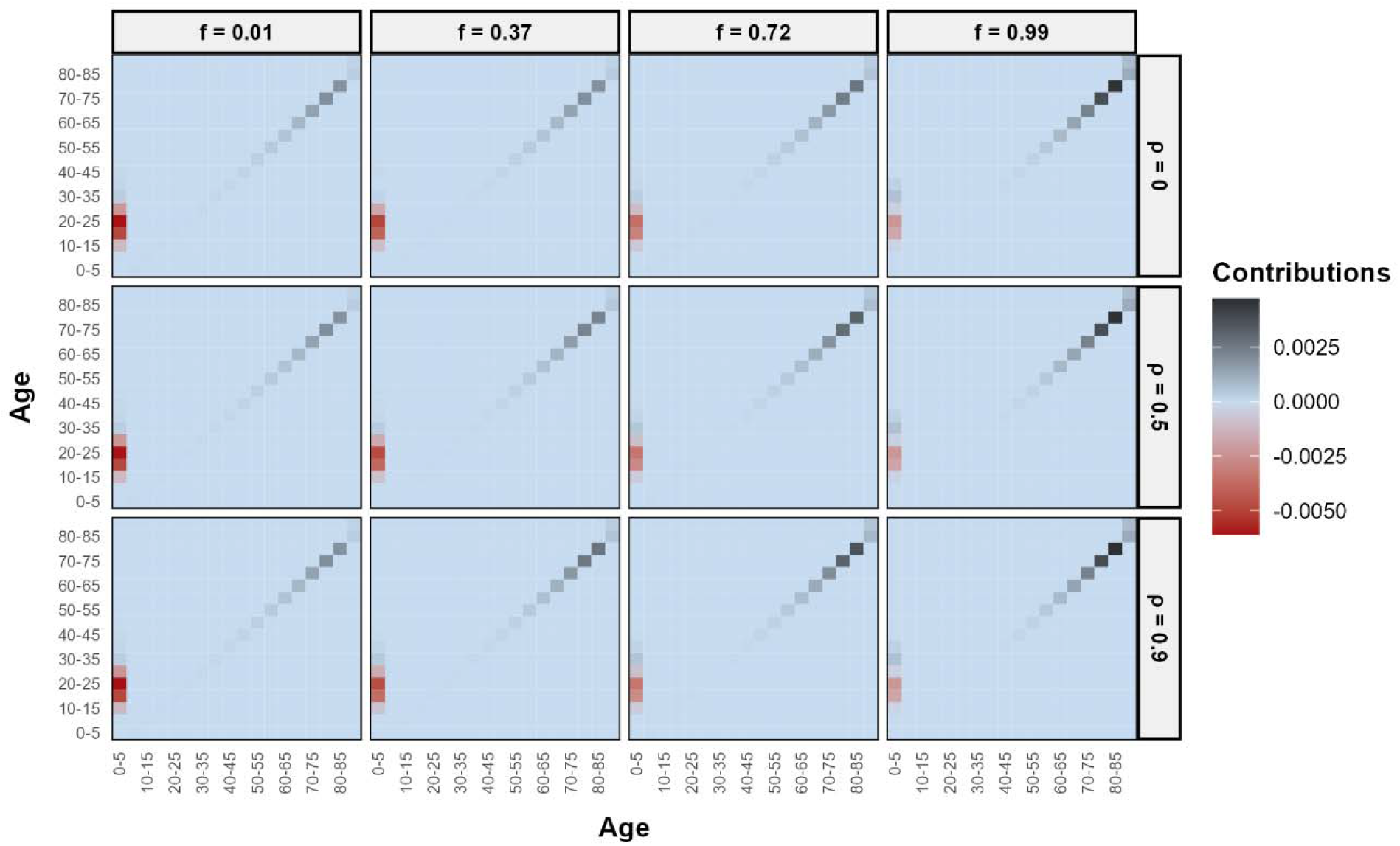
Lower adult survival rates especially at ages more than 60 years for treatment 3 *(10-50%, galloping inflation)* in Italy contributes positively to the differences in the stochastic growth rate between treatment 2 *(3-10%, walking inflation)* and 3 at all stationary frequency *(f)* and temporal autocorrelation *(ρ)*. The contribution of adult survival to the growth difference increases with increasing frequency. However, higher fertility at early reproductive ages (20-25 yrs) contributes negatively to the differences in stochastic growth rate between treatment 2 and 3. Matrix plot shows the contribution of the matrix elements **A**_***i***_, integrated over all environmental states, to the difference in stochastic growth rate log λ_s_ of treatment 2 *(3-10%, walking inflation)* and treatment 3 *(10-50%, galloping inflation)*, for Italy. Growth rate and sensitivities are calculated by Markov chain simulation with T = 1000 iterations.

**Fig. S10.**
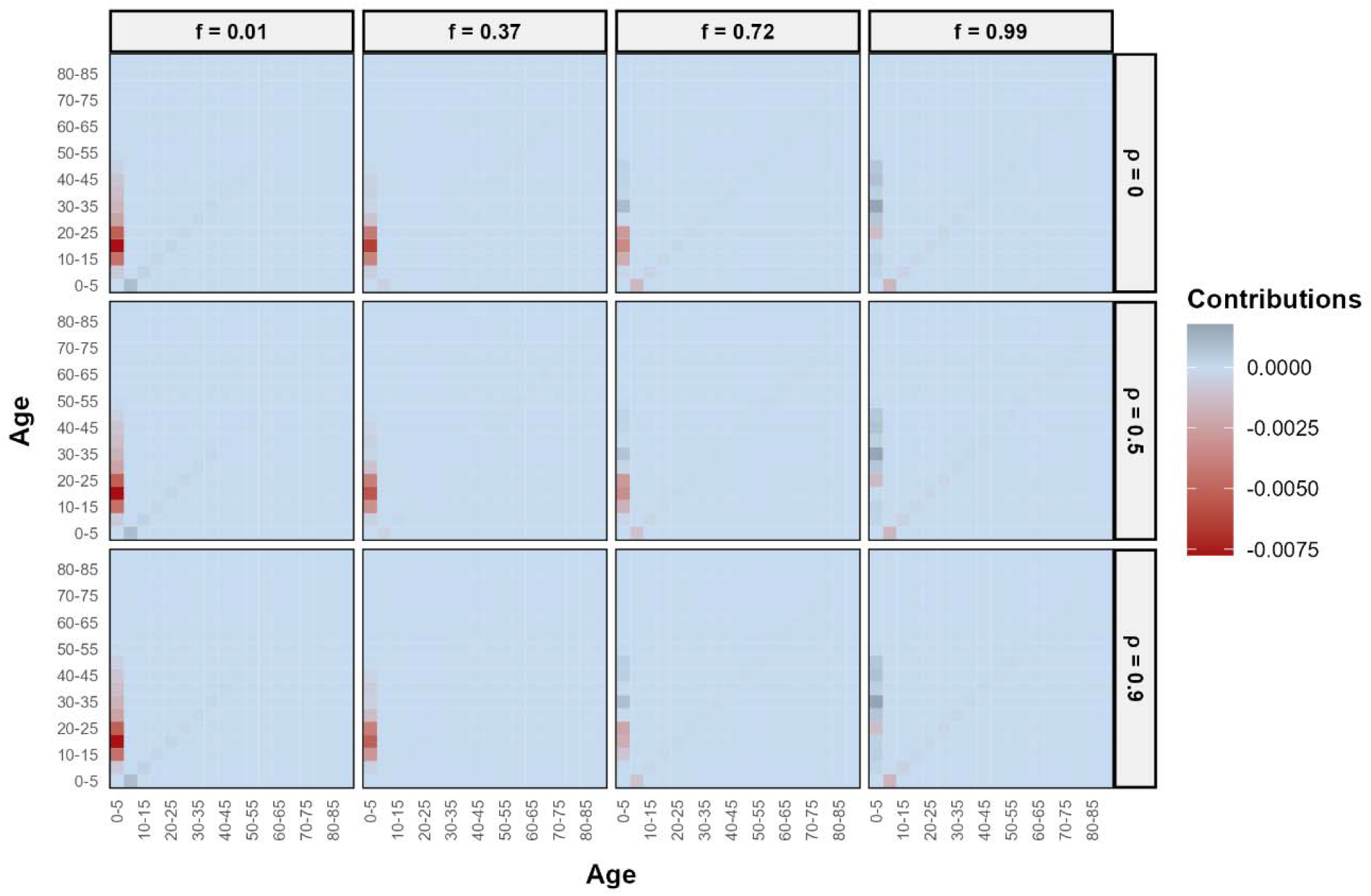
Lower fertility rates at early reproductive ages (10-25 years) for treatment 4 (>50%, *hyperinflation)* in Sudan contributes negatively to the differences in stochastic growth rate between treatment 3 *(10-50%, galloping inflation)* and treatment 4 at all stationary frequency (f) and temporal autocorrelation (p). Matrix plot shows the contribution of the **A**_***i***_, integ rated over all environmental states, to the difference in stochastic growth rate log λ_s_ of treatment 3 *(10-50%, galloping inflation)* and treatment 4 (>50%, *hyperinflation)*, for Sudan. Growth rate and sensitivities are calculated by Markov chain simulation with T = 1000 iterations.

**Fig. S11.**
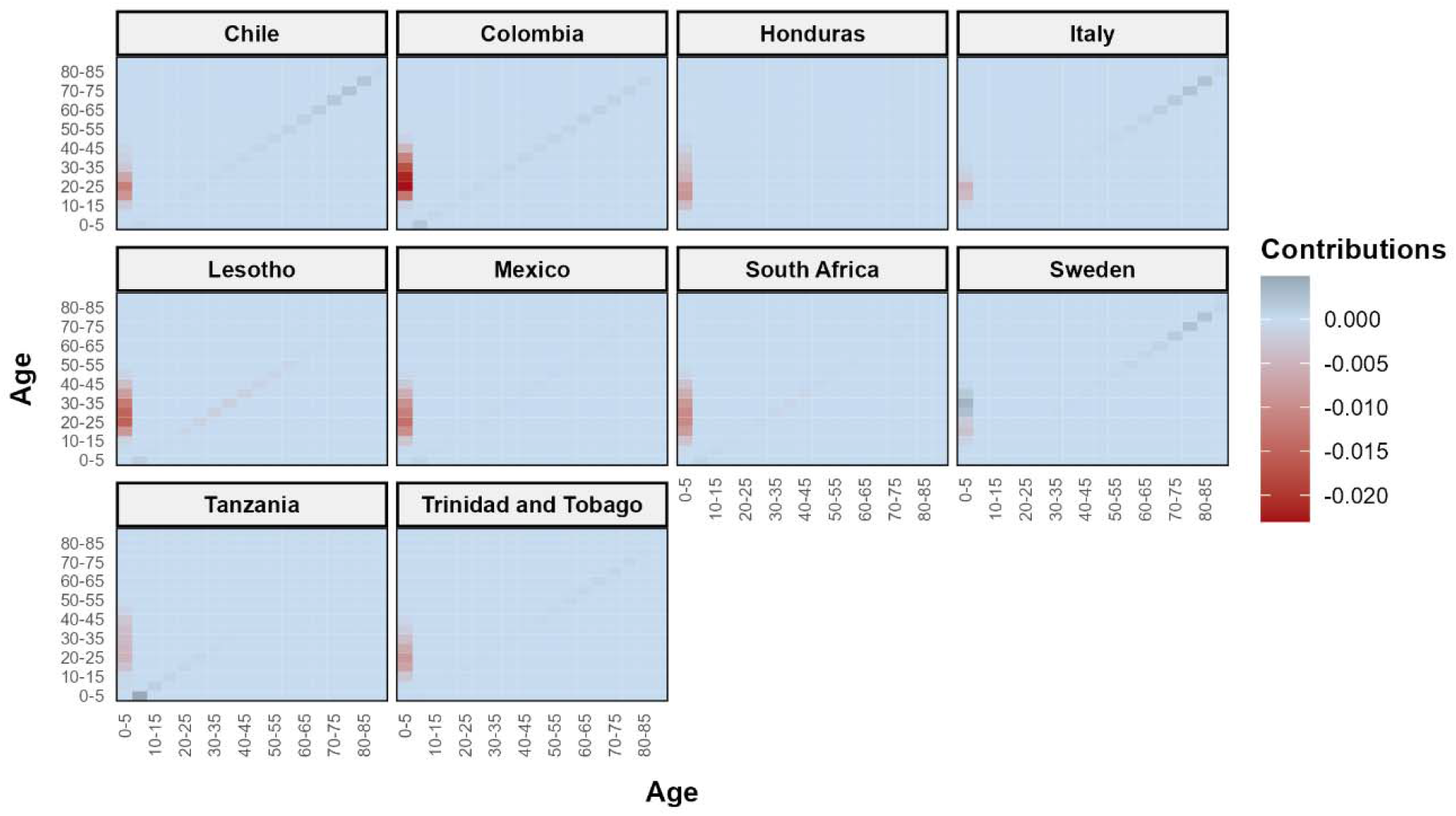
Disadvantage in fertility rates at all ages especially 15-40 years for treatment 3 *(10-50%, galloping inflation)* contributes negatively to the growth disadvantage of treatment 3 over treatment 2 *(3-10%, walking inflation)* in most of the countries. The contribution of differences in fertility is high in countries such as Colombia, Lesotho, and Mexico, shown in red colour. However, in Sweden, fertility at late reproductive years (>30 years) contributes positively to the growth rate differences between treatment 3 and 4. The differenes in adult survival contributes positively to the growth rate difference in most of the countries, such as Chile, Colombia, Italy, and Sweden, shown in the grey colour. However, in countries such as Colombia and Tanzania, juvenile survival contributes positively to the growth rate differences. Matrix plot shows the contribution of the **A**_***i***_, integrated over all environmental states, to the difference in stochastic growth rate log λ_s_ of treatment 2 (*3-10%, walking inflation)* and treatment 3 (*10-50%, galloping inflation)*, for select countries. Growth rates and sensitivities are calculated by Markov chain simulation with *T* = 1000 iterations.

**Fig. S12.**
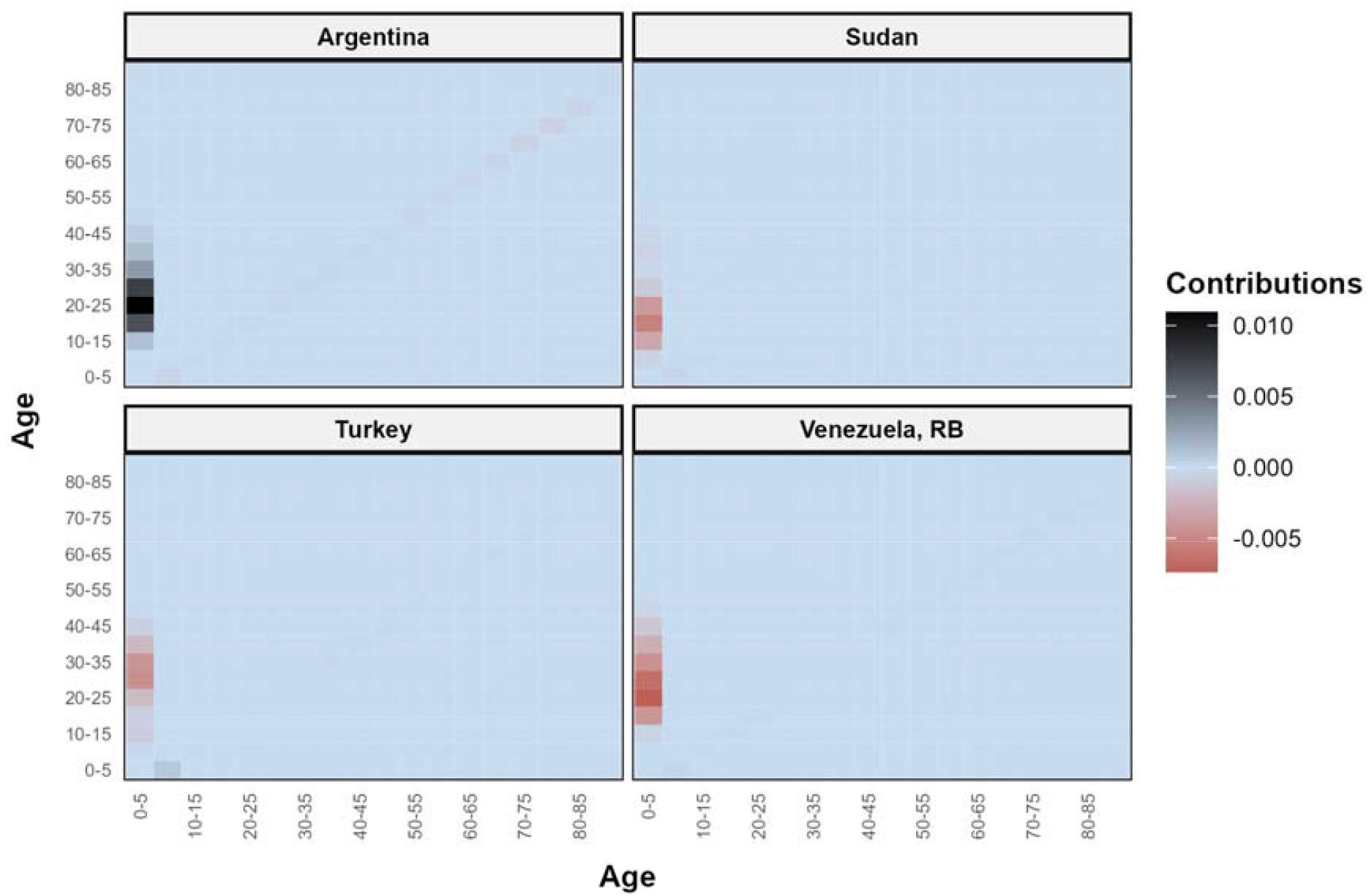
Differences in fertility rates at all ages, especially 20-30 years contributes negatively to the differences between treatment 4 *(>50%, hyperinflation)* and treatment 3 *(10-50%, galloping inflation)* in all countries, except in Argentina. The negative contribution of the fertility rates are shown in the red colour. In Argentina, the differences in fertility rates at treatment 3 and treatment 4 contributes positively to the treatment effect on stochastic growth rate, shown in the grey colour. Moreover, the differences in juvenile survival contributes positively to the growth rate differences between treatment 3 and 4 in Turkey. Matrix plot shows the contribution of the **A**_***i***_, integrated over all environmental states, to the difference in stochastic growth rate log λ_s_ of treatment 3 (*10-50%, galloping inflation)* and treatment 4 *(>50%, hyperinflation)*, for select countries. Growth rates and sensitivities are calculated by Markov chain simulation with *T* = 1000 iterations.

